# Epigenetic reprogramming of a distal developmental enhancer cluster drives *SOX2* overexpression in breast and lung cancer

**DOI:** 10.1101/2023.03.14.532258

**Authors:** Luis E. Abatti, Patricia Lado-Fernández, Linh Huynh, Manuel Collado, Michael M. Hoffman, Jennifer A. Mitchell

## Abstract

Enhancer reprogramming has been proposed as a key source of transcriptional dysregulation during tumorigenesis, but the molecular mechanisms underlying this process remain unclear. Here, we identify an enhancer cluster required for normal development that is aberrantly activated in breast and lung carcinoma. Deletion of the SRR124–134 cluster disrupts transcription of the *SOX2* oncogene and dysregulates genome-wide chromatin accessibility in cancer cells. Analysis of primary tumors reveals a correlation between chromatin accessibility at this cluster and *SOX2* overexpression in breast and lung cancer patients. We demonstrate that FOXA1 is an activator and NFIB is a repressor of SRR124–134 activity and *SOX2* transcription in cancer cells, revealing a co-opting of the regulatory mechanisms involved in early development. Notably, we show that the conserved SRR124 and SRR134 regions are essential during mouse development, where homozygous deletion results in the lethal failure of esophageal-tracheal separation. These findings provide insights into how developmental enhancers can be reprogrammed during tumorigenesis and underscore the importance of understanding enhancer dynamics during development and disease.

## INTRODUCTION

Developmental enhancers are commissioned during early embryogenesis, as transcription factors progressively restrict the epigenome through the repression of regulatory regions associated with pluripotency (Zhu et al. 2013; Hawkins et al. 2010) and the activation of enhancers that control the expression of lineage-specific developmental genes (Rada-Iglesias et al. 2011; Creyghton et al. 2010; Rubin et al. 2017). This establishes a cell type-specific epigenetic regulatory “memory” that maintains cell lineage commitment and reinforces transcriptional programs (Stergachis et al. 2013). As cells mature and development ends, developmental-associated enhancers are decommissioned, and the enhancer landscape becomes highly restrictive and developmentally stable (Stergachis et al. 2013). This landscape, however, becomes profoundly disturbed during tumorigenesis, as cancer cells aberrantly acquire euchromatin features at regions near oncogenes (Hnisz et al. 2013; Lovén et al. 2013) that are often associated with earlier stages of cell lineage specification (Stergachis et al. 2013). This “enhancer reprogramming” has been proposed to result in a dysfunctional state that causes widespread abnormal gene expression and cellular plasticity (Fu et al. 2019; Roe et al. 2017; Bi et al. 2020; Pomerantz et al. 2015; Lupien et al. 2008). Although the misactivation of enhancers has been suggested as a major source of transcriptional dysregulation (reviewed in Richart et al. 2021; Okabe and Kaneda 2021), it remains largely unclear how this mechanism unfolds during the progression of cancer. To study this process, we evaluated cis-regulatory elements involved in driving transcription during normal development and disease.

SRY-box transcription factor 2 (SOX2) is a pioneer transcription factor required for pluripotency maintenance in embryonic stem cells (Avilion 2003; Chew et al. 2005), involved in reprogramming differentiated cells to induced pluripotent stem cells in mammals (Takahashi and Yamanaka 2006; Takahashi et al. 2007; Yu et al. 2007), and acts as an oncogene in several different types of cancer (reviewed in Wuebben and Rizzino 2017; Novak et al. 2019). During later development, SOX2 is also required for tissue morphogenesis and homeostasis of the brain (Graham et al. 2003), eyes (Taranova et al. 2006), esophagus (Que et al. 2007), inner ear (Kiernan et al. 2005), lungs (Gontan et al. 2008), skin (Driskell et al. 2009), stomach (Francis et al. 2019), taste buds (Okubo et al. 2006) and trachea (Que et al. 2009) in both human and mouse. In these tissues, *SOX2* expression is regulated precisely in space and time at critical stages of development, although in most cases the cis-regulatory regions that mediate this precision remain unknown. For example, proper levels of *Sox2* expression are required for the complete separation of the anterior foregut into the esophagus and trachea in mice (Que et al. 2007; Teramoto et al. 2020; Chakraborty et al. 2023) and in humans (Zenteno et al. 2006; Williamson et al. 2006; Chassaing et al. 2007), as the disruption of *Sox2* expression leads to an abnormal developmental condition known as esophageal atresia with distal tracheoesophageal fistula (EA/TEF) (reviewed in Brunner and van Bokhoven 2005; Que et al. 2006). After the anterior foregut is properly separated, *Sox2* expression ranges from the esophagus to the stomach in the gut (Que et al. 2007; Francis et al. 2019), and throughout the trachea, bronchi, and upper portion of the lungs in the developing airways (Que et al. 2009). Proper branching morphogenesis at the tip of the lungs, however, requires temporary downregulation of *Sox2*, followed by reactivation after lung bud establishment (Gontan et al. 2008). *Sox2* also retains an essential function in multiple mature epithelial tissues, where it is highly expressed in proliferative and self-renewing adult stem cells necessary for maintaining and replacing terminally differentiated cells within the epithelium of the brain, bronchi, esophagus, stomach, and trachea (Arnold et al. 2011; Que et al. 2009; Tompkins et al. 2009; Francis et al. 2019). However, the expression of *SOX2* becomes repressed within terminally differentiated epithelial cells in these tissues (Arnold et al. 2011).

As an oncogene, overexpression of *SOX2* is linked to increased cellular replication rates, aggressive tumor grades, and poor patient outcomes in breast carcinoma (BRCA) (Chen et al. 2008; Leis et al. 2012; Liu et al. 2018b; Meng et al. 2020; Piva et al. 2014); colon adenocarcinoma (COAD) (Takeda et al. 2018; Talebi et al. 2015; Zhang et al. 2020b; Zhu et al. 2021); glioblastoma (GBM) (Alonso et al. 2011; Cox et al. 2012; Gangemi et al. 2009; Hägerstrand et al. 2011); liver hepatocellular carcinoma (LIHC) (Sun et al. 2013); lung adenocarcinoma (LUAD) (Chou et al. 2013; Sholl et al. 2010; Nakatsugawa et al. 2011); and lung squamous cell carcinoma (LUSC) (Bass et al. 2009; Hussenet et al. 2010). These clinical and molecular characteristics arise from the participation of *SOX2* in the formation and maintenance of tumor-initiating cells that resemble tissue progenitor cells, as evidenced by BRCA (Domenici et al. 2019; Piva et al. 2014; Simões et al. 2011), GBM (Bulstrode et al. 2017; Gangemi et al. 2009; Jeon et al. 2011; Zhang et al. 2020a, 24), LUAD (Singh et al. 2012), and LUSC (Boumahdi et al. 2014) studies. *SOX2* knockdown, on the other hand, often results in diminished levels of cell replication, invasion, and treatment resistance in these tumor types (Bass et al. 2009; Berezovsky et al. 2014; Chen et al. 2008; Chou et al. 2013; Fang et al. 2011; Nakatsugawa et al. 2011; Piva et al. 2014; Stolzenburg et al. 2012). Despite the involvement of *SOX2* in the progression of multiple types of cancer, little is known about the mechanisms that cause *SOX2* overexpression during tumorigenesis. Two proximal enhancers were once deemed crucial for driving *Sox2* expression during early development: *<u>S</u>ox2* <u>R</u>egulatory <u>R</u>egion 1 (SRR1) and SRR2 (Ferri et al. 2004; Tomioka et al. 2002; Zappone et al. 2000). Deletion of SRR1 and SRR2, however, has no effect on *Sox2* expression in mouse embryonic stem cells (Zhou et al. 2014a). In contrast, deletion of a distal *<u>S</u>ox2* <u>C</u>ontrol <u>R</u>egion (SCR), 106 kb downstream of the *Sox2* promoter, causes a profound loss of *Sox2* expression in mouse embryonic stem cells (Li et al. 2014; Zhou et al. 2014a) and in blastocysts, where SCR deletion causes peri-implantation lethality (Chakraborty et al. 2023). The contribution of these regulatory regions in driving *SOX2* expression during tumorigenesis, however, remains poorly defined.

Here, we investigated the mechanisms underlying *SOX2* overexpression in cancer. We found that, in breast and lung cancer, *SOX2* is driven by a novel developmental enhancer cluster we termed SRR124–134, rather than the previously identified SRR1, SRR2, or the SCR. This novel distal cluster contains two regions located 124 and 134 kb downstream of the *SOX2* promoter that drive transcription in breast and lung cancer cells. Deletion of this cluster results in significant *SOX2* downregulation, leading to genome-wide changes in chromatin accessibility and a globally disrupted transcriptome. The SRR124–134 cluster is highly accessible in most breast and lung patient tumors, where chromatin accessibility at these regions is correlated with *SOX2* overexpression and is regulated positively by FOXA1 and negatively by NFIB. Finally, we found that both SRR124 and SRR134 are highly conserved in the mouse and are essential for postnatal survival, as homozygous deletion of their homologous regions results in lethal EA/TEF. These findings serve as a prime example of how different types of cancer cells reprogram enhancers that were decommissioned during development to drive the expression of oncogenes during tumorigenesis.

## RESULTS

### Two regions downstream of *SOX2* gain enhancer features in cancer cells

*SOX2* overexpression occurs in multiple types of cancer (reviewed in Wuebben and Rizzino 2017; Novak et al. 2019). To examine which cancer types have the highest levels of *SOX2* upregulation, we performed differential expression analysis by calculating the log_2_ fold change (log_2_ FC) of *SOX2* transcription from 21 TCGA primary solid tumors (see Supplementary Table S1 for cancer type abbreviations) compared to normal tissue samples (Hoadley et al. 2014). We found that BRCA (log_2_ FC = 3.31), COAD (log_2_ FC = 1.38), GBM (log_2_ FC = 2.05), LIHC (log_2_ FC = 3.22), LUAD (log_2_ FC = 1.36), and LUSC (log_2_ FC = 4.91) tumors had the greatest *SOX2* upregulation (log_2_ FC > 1; FDR-adjusted *Q* < 0.01; Figure 1A, Supplementary Table S2). As a negative control, we ran this same analysis using the housekeeping gene *PUM1* (Krasnov et al. 2019) and found no cancer types with significant upregulation of this gene (Supplementary Figure S1A, Supplementary Table S3).

**Figure 1:**
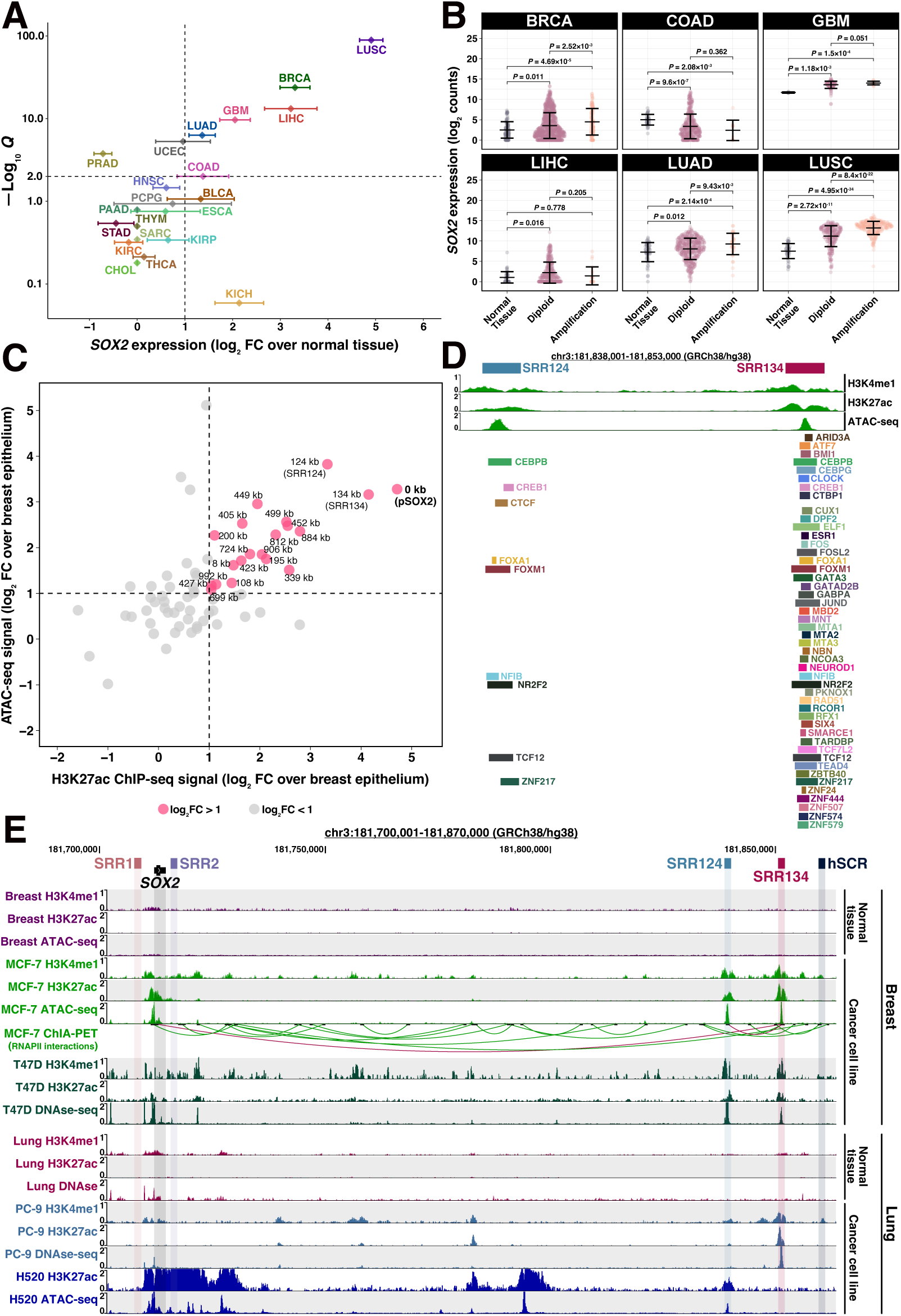
A cluster 124–134 kilobases downstream of *SOX2* gains enhancer features in cancer cells. **(A)** Super-logarithmic RNA-seq volcano plot of *SOX2* expression from 21 cancer types compared to normal tissue (Hoadley et al. 2014). Cancer types with log_2_ FC > 1 and FDR-adjusted *Q* < 0.01 were considered to significantly overexpress *SOX2*. Error bars: standard deviation. **(B)** *SOX2* log_2_-normalized expression (log_2_ counts) associated with the *SOX2* copy number from BRCA (n = 1174), COAD (n = 483), GBM (n = 155), LIHC (n = 414), LUAD (n = 552), and LUSC (n = 546) patient tumors (Hoadley et al. 2014). RNA-seq reads were normalized to library size using DESeq2 (Love et al. 2014). Error bars: standard deviation. Significance analysis by Dunn’s test (Dunn 1964) with Holm correction (Holm 1979). **(C)** 1,500 bp genomic regions within ± 1 Mb from the *SOX2* transcription start site (TSS) that gained enhancer features in MCF-7 cells (ENCODE Project Consortium 2012) compared to normal breast epithelium (Roadmap Epigenomics Consortium et al. 2015). Regions that gained both ATAC-seq and H3K27ac ChIP-seq signal above our threshold (log_2_ FC > 1, dashed line) are highlighted in pink. Each region was labelled according to their distance in kilobases (kb) to the *SOX2* promoter (pSOX2, bolded). **(D)** ChIP-seq signal for H3K4me1 and H3K27ac, ATAC-seq signal, and transcription factor ChIP-seq peaks at the SRR124–134 cluster in MCF-7 cells. Datasets from ENCODE (ENCODE Project Consortium 2012)**. (E)** UCSC Genome Browser (Kent et al. 2002) display of H3K4me1 and H3K27ac ChIP-seq signal, DNAse-seq and ATAC-seq chromatin accessibility signal, and ChIA-PET RNA Polymerase II (RNAPII) interactions around the *SOX2* gene within breast (normal tissue and 2 BRCA cancer cell lines) and lung (normal tissue, one LUAD, and one LUSC cancer cell lines) samples (ENCODE Project Consortium 2012; Chan et al. 2018; Sato et al. 2019; Gopi and Kidder 2021). Relevant RNAPII interactions (between SRR124 and SRR134, and between SRR134 and pSOX2) are highlighted in maroon.

Next, we divided BRCA, COAD, GBM, LIHC, LUAD, and LUSC patients (n = 3,064) into four groups according to their *SOX2* expression. Gene expression levels were measured by RNA-seq counts normalized by library size and transformed to a log_2_ scale, hereinafter referred to as log_2_ counts. Cancer patients within the top group (25% highest *SOX2* expression; log_2_ counts > 10.06) have a significantly (*P* = 1.27×10^-23^, log-rank test) lower overall probability of survival compared to cancer patients within the bottom group (25% lowest *SOX2* expression; log_2_ counts < 1.68) (Supplementary Figure S1B, Supplementary Table S4). We also examined the relationship between *SOX2* copy number and *SOX2* overexpression within these six tumor types. Although previous studies have shown that *SOX2* is frequently amplified in squamous cell carcinoma (Bass et al. 2009; Hussenet et al. 2010; Maier et al. 2011; Liu et al. 2021), we found that most BRCA (88%), COAD (98%), GBM (91%), LIHC (94%), and LUAD (92%) tumors were diploid for *SOX2*. In addition, BRCA (*P* = 0.011, Holm-adjusted Dunn’s test), GBM (*P* = 1.18×10^-3^), LIHC (*P* = 0.016), LUAD (*P* = 0.012), and LUSC (*P* = 2.72×10^-11^) diploid tumors significantly overexpressed *SOX2* compared to normal tissue (Figure 1B, Supplementary Table S5). This indicates that gene amplification is dispensable for driving *SOX2* overexpression in most cancer types.

We investigated whether the *SOX2* locus gains epigenetic features associated with active enhancers in cancer cells. Enhancer features commonly include accessible chromatin determined by either Assay for Transposase Accessible Chromatin with high-throughput sequencing (ATAC-seq) (Buenrostro et al. 2013) or DNase I hypersensitive sites sequencing (DNase-seq) (Boyle et al. 2008), and histone modifications including histone H3 lysine 4 monomethylation (H3K4me1) and histone H3 lysine 27 acetylation (H3K27ac) (Heintzman et al. 2007, 2009). To study gains in enhancer features within the *SOX2* locus, we initially focused our analyses on luminal A breast cancer, the most common subtype of BRCA to significantly (*P* = 0.021, Tukey’s test) overexpress *SOX2* (Supplementary Figure S1C) (Hoadley et al. 2014; Berger et al. 2018). MCF-7 cells are a widely used ER^+^/PR^+^/HER2^-^ luminal A breast adenocarcinoma model (Soule et al. 1973), which have been previously described to overexpress *SOX2* (Chen et al. 2008; Liang et al. 2013; Ling et al. 2012; Stolzenburg et al. 2012). After confirming that *SOX2* is one of the most upregulated genes in MCF-7 cells (log_2_ FC = 10.75; FDR-adjusted *Q* = 2.20×10^-36^; Supplementary Figure S1D, Supplementary Table S6) compared to normal breast epithelium (Roadmap Epigenomics Consortium et al. 2015), we contrasted their chromatin accessibility and histone modifications (ENCODE Project Consortium 2012). By intersecting 1,500 bp regions that contain at least 500 bp overlap between H3K27ac and ATAC-seq peaks, we found that 19 putative enhancers gained (log_2_ FC > 1) both these features within ± 1 Mb from the *SOX2* transcription start site (TSS) in MCF-7 cells (Figure 1C, Supplementary Table S7). Besides the SOX2 promoter (pSOX2), we identified a downstream cluster containing two regions that have gained the highest ATAC-seq and H3K27ac signal in MCF-7 cells: SRR124 (124 kb downstream of pSOX2) and SRR134 (134 kb downstream of pSOX2). The previously described SRR1, SRR2 (Ferri et al. 2004; Tomioka et al. 2002; Zappone et al. 2000), and the human ortholog of the mouse SCR (hSCR) (Li et al. 2014; Zhou et al. 2014a), however, lacked substantial gains in enhancer features within MCF-7 cells.

Alongside gains in chromatin features, another characteristic of active enhancers is the binding of numerous (> 10) transcription factors (Chen et al. 2012; Mitchell et al. 2012; Singh et al. 2021). Chromatin Immunoprecipitation Sequencing (ChIP-seq) data from ENCODE (ENCODE Project Consortium 2012) on 117 transcription factors revealed 48 different factors present at the SRR124–134 cluster in MCF-7 cells, with the majority (47) of these factors present at SRR134 (Figure 1D). Transcription factors bound at both SRR124 and SRR134 include CEBPB, CREB1, FOXA1, FOXM1, NFIB, NR2F2, TCF12, and ZNF217. An additional feature of distal enhancers is that they contact their target genes through long-range chromatin interactions (Tolhuis et al. 2002; Carter et al. 2002). Chromatin Interaction Analysis by Paired-End-Tag sequencing (ChIA-PET) (Fullwood et al. 2009) showed two interesting RNA Polymerase II (RNAPII)-mediated chromatin interactions in MCF-7 cells: one between the *SOX2* gene and SRR134, and one between SRR124 and SRR134 (Figure 1E). Beyond MCF-7 cells, we found that H520 (LUSC), PC-9 (LUAD), and T47D (luminal A BRCA) cancer cell lines, which display varying levels of *SOX2* expression (Supplementary Figure S1E), also gained substantial enhancer features at SRR124 and SRR134 when compared to normal tissue (Figure 1E) (ENCODE Project Consortium 2012; Chan et al. 2018; Sato et al. 2019; Gopi and Kidder 2021). Together, these data suggest SRR124 and SRR134 could be active enhancers driving *SOX2* transcription in BRCA, LUAD, and LUSC.

### The SRR124–134 cluster is essential for *SOX2* expression in BRCA and LUAD cells

To assess SRR124 and SRR134 enhancer activity alongside the embryonic-associated SRR1, SRR2, and hSCR regions, we used a reporter vector containing the firefly luciferase gene under the control of a minimal promoter (minP, pGL4.23). We transfected each enhancer construct into the BRCA (MCF-7, T47D), LUAD (PC-9), and LUSC (H520) cell lines and measured luciferase activity as a relative fold change (FC) compared to the empty minP vector. SRR134 demonstrated the strongest enhancer activity, with the MCF-7 (FC = 6.42; *P* < 2×10^-16^, Dunnett’s test), T47D (FC = 3.36; *P* = 9.34×10^-10^), H520 (FC = 2.37; *P* = 1.22×10^-6^), and PC-9 (FC = 2.03; *P* = 9.79×10^-5^) cell lines displaying a significant increase in luciferase activity compared to minP (Figure 2A). SRR124 also showed a modest, significant increase in luciferase activity compared to minP in the MCF-7 (FC = 1.53; *P* = 4.27×10^-2^), T47D (FC = 1.80; *P* = 4.57×10^-2^), and PC-9 (FC = 1.60; *P* = 4.27×10^-2^) cell lines. The SRR1, SRR2, and hSCR enhancers, however, showed no significant enhancer activity (*P* > 0.05) in all four cell lines.

**Figure 2:**
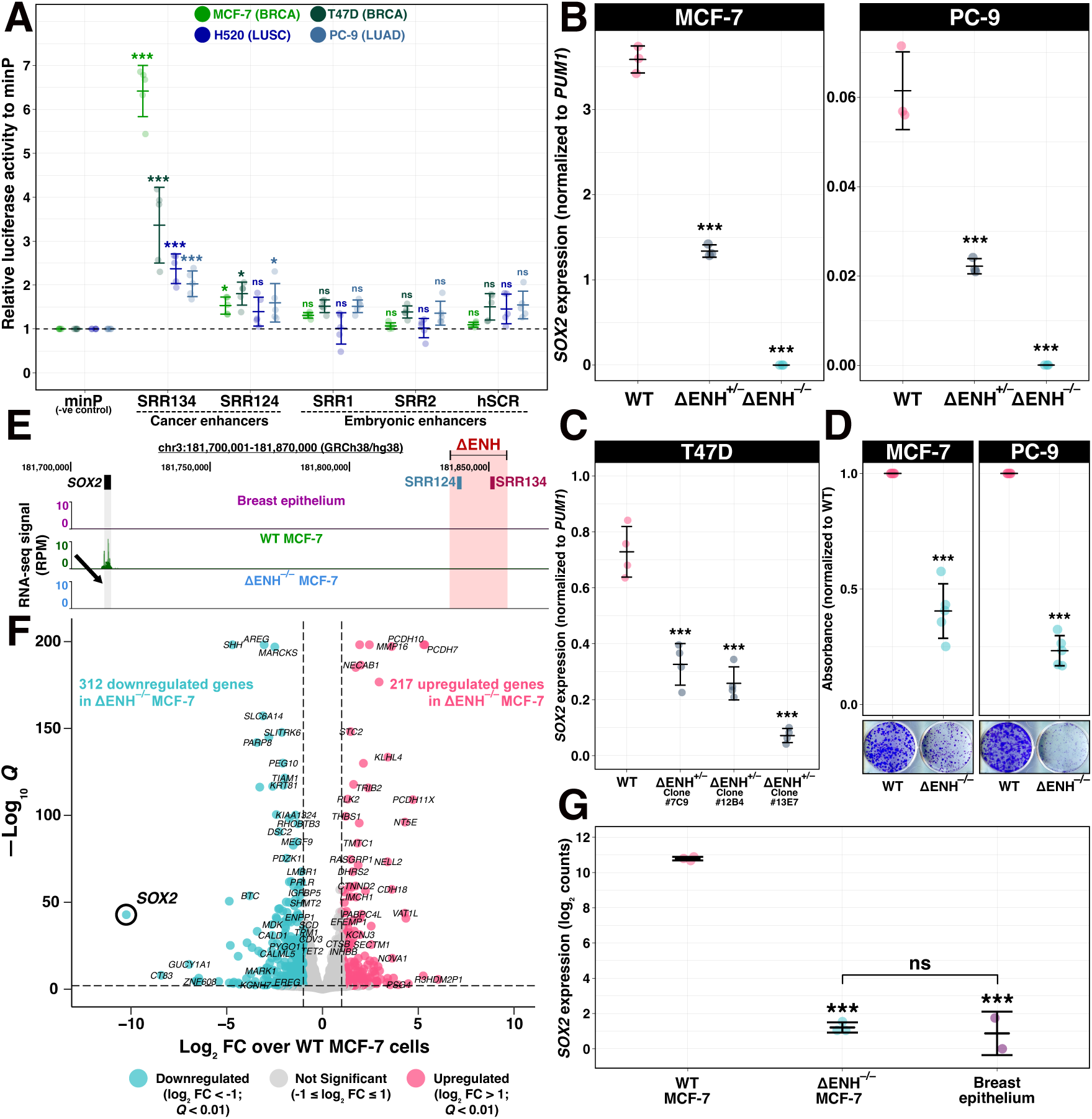
The SRR124–134 cluster drives *SOX2* overexpression in MCF-7, PC-9, and T47D cells. **(A)** Enhancer reporter assay comparing luciferase activity driven by the SRR1, SRR2, SRR124, SRR134, and hSCR regions to an empty vector containing only a minimal promoter (minP). Enhancer constructs were assayed in the BRCA (MCF-7, T47D), LUAD (PC-9), and LUSC (H520) cell lines. Dashed line: average activity of minP. Error bars: standard deviation. Significance analysis by Dunnett’s test (n = 5; * *P* < 0.05, *** *P* < 0.001, ns: not significant) (Dunnett 1955). **(B)** RT-qPCR analysis of *SOX2* transcript levels in SRR124–134 heterozygous- (ΔENH^+/–^) and homozygous- (ΔENH^−/−^) deleted MCF-7 (BRCA) and PC-9 (LUAD) clones compared to WT cells. Error bars: standard deviation. Significance analysis by Dunnett’s test (n = 3; *** *P* < 0.001). **(C)** RT-qPCR analysis of *SOX2* transcript levels in three independent SRR124–134 heterozygous-deleted (ΔENH^+/–^) T47D clonal isolates compared to WT cells. Error bars: standard deviation. Significance analysis by Dunnett’s test (n = 4; *** *P* < 0.001). **(D)** Crystal violet absorbance (570 nm) from a colony formation assay with WT and ΔENH^−/−^ MCF-7 and PC-9 cells. Total absorbance was normalized to the average absorbance from WT cells within each cell line. Significance analysis by t-test with Holm correction (n = 5; *** *P* < 0.001). **(E)** UCSC Genome Browser (Kent et al. 2002) view of the SRR124–134 cluster deletion in ΔENH^−/−^ MCF-7 cells with RNA-seq tracks from normal breast epithelium (Roadmap Epigenomics Consortium et al. 2015), WT and ΔENH^−/−^ MCF-7 cells. Arrow: reduction in RNA-seq signal at the *SOX2* gene in ΔENH^−/−^ MCF-7 cells. **(F)** Volcano plot with DESeq2 (Love et al. 2014, 2) differential expression analysis between ΔENH^−/−^ and WT MCF-7 cells. Blue: 312 genes that significantly lost expression (log_2_ FC < -1; FDR-adjusted *Q* < 0.01) in ΔENH^−/−^ MCF-7 cells. Pink: 217 genes that significantly gained expression (log_2_ FC > 1; *Q* < 0.01) in ΔENH^−/−^ MCF-7 cells. Grey: 35,891 genes that maintained similar (-1 ≤ log_2_ FC ≤ 1) expression between ΔENH^−/−^ and WT MCF-7 cells. **(G)** Comparison of *SOX2* transcript levels between WT MCF-7 and either ΔENH^−/−^ MCF-7 or normal breast epithelium cells (Roadmap Epigenomics Consortium et al. 2015), and between ΔENH^−/−^ MCF-7 and normal breast epithelium cells. RNA-seq reads were normalized to library size using DESeq2 (Love et al. 2014). Error bars: standard deviation. Significance analysis by Tukey’s test (*** *P* < 0.001, ns: not significant) (Tukey 1949).

Although reporter assays can be used to assess enhancer activity, enhancer knockout approaches remain the current gold standard method for enhancer validation (Moorthy et al. 2017; Tobias et al. 2021). To investigate whether the SRR124–134 cluster drives *SOX2* expression in cancer cells, we used CRISPR/Cas9 to delete this cluster from breast (MCF-7, T47D) and lung (H520, PC-9) cancer cell lines. RT-qPCR showed that homozygous SRR124–134 deletion (ΔENH^−/−^) causes a profound (> 99.5%) and significant (*P* < 0.001, Dunnett’s test) loss of *SOX2* expression in both the MCF-7 and PC-9 cell lines (Figure 2B). Immunoblot analysis confirmed the depletion of the SOX2 protein in ΔENH^−/−^ MCF-7 cells (Supplementary Figure S2A). Heterozygous SRR124– 134 deletion (ΔENH^+/–^) also significantly (*P* < 0.001) reduced *SOX2* expression by ∼60% in both MCF-7 and PC-9 cells (Figure 2B). Although we were unable to isolate a homozygous deletion clone from T47D cells, multiple independent heterozygous ΔENH^+/–^ T47D clonal isolates also showed a significant downregulation (>50%; *P* < 0.001) in *SOX2* expression (Figure 2C). Interestingly, we did not find a significant (*P* > 0.05) impact on *SOX2* expression in ΔENH^+/–^ or ΔENH^−/−^ H520 cells (Supplementary Figure S2B), which indicates that *SOX2* transcription is sustained by a different mechanism in these cells. To assess the impact of the loss of *SOX2* expression in the tumor initiation capacity of enhancer-deleted cells, we performed a colony formation assay with MCF-7 and PC-9 ΔENH^−/−^ cells. We found that both MCF-7 (*P* = 3.53×10^-4^, t-test) and PC-9 (*P* = 1.26×10^-5^) ΔENH^−/−^ cells showed a significant decrease (> 50%) in their ability to form colonies compared to WT cells (Figure 2D), further indicating that SRR124–134-driven *SOX2* overexpression is required to sustain high tumor initiation capacity in BRCA and LUAD.

Next, we performed total RNA sequencing (RNA-seq) to measure changes in the transcriptome of ΔENH^−/−^ MCF-7 cells compared to WT MCF-7 cells. As expected, all three replicates of each genotype clustered together (Supplementary Figure S2C). In addition to *SOX2* downregulation (Figure 2E), differential expression analysis showed a total of 529 genes differentially (|log_2_ FC| > 1; FDR-adjusted *Q* < 0.01) expressed in ΔENH^−/−^ MCF-7 cells (Figure 2F, Supplementary Table S8). From these, 312 genes significantly lost expression (59%), whereas 217 (41%) genes significantly gained expression in ΔENH^−/−^ MCF-7 cells compared to WT MCF-7 cells (Supplementary Figure S2D). *SOX2* was the gene with the highest loss in expression (log_2_ FC = -10.24; *Q* = 1.23×10^-43^) in ΔENH^−/−^ MCF-7 cells, followed by *CT83* (log_2_ FC = -8.43; *Q* = 1.07×10^-8^), and *GUCY1A1* (log_2_ FC = -6.96; *Q* = 5.09×10^-15^). On the other hand, genes with the most significant gain in expression within ΔENH^−/−^ MCF-7 cells included the protocadherins *PCDH7* (log_2_ FC = 5.34; *Q* < 1×10^-200^), *PCDH10* (log_2_ FC = 5.29; *Q* < 1×10^-200^), and *PCDH11X* (log_2_ FC = 4.73; Q = 9.29×10^-110^). In addition, deletion of the SRR124–134 cluster reduced *SOX2* expression back to the levels found in normal breast epithelium (*P* = 0.48, Tukey’s test) (ENCODE Project Consortium 2012; Roadmap Epigenomics Consortium et al. 2015) (Figure 2G). Together, these data confirm that the SRR124–134 cluster drives *SOX2* overexpression in BRCA and LUAD.

### SOX2 regulates pathways associated with epithelium development in luminal A BRCA

Because SOX2 regulates cell proliferation and differentiation pathways in other epithelial cells (Tompkins et al. 2009, 2011), we decided to further investigate the molecular function of SOX2 in luminal A BRCA cells by utilizing our *SOX2*-depleted ΔENH^−/−^ MCF-7 cell model. Gene Set Enrichment Analysis (GSEA) showed a significant (FDR-adjusted *Q* < 0.05) depletion of multiple epithelium-associated processes within the transcriptome of ΔENH^−/−^ MCF-7 cells, as indicated by the normalized enrichment score [NES] < 1 (Supplementary Table S9). These processes included epidermis development (NES = -1.93; *Q* = 0.001; Figure 3A), epithelial cell differentiation (NES = -1.67; *Q* = 0.007; Figure 3B), and cornification (NES = -2.11; *Q* = 0.006; Figure 3C). This suggests that SOX2 regulates epithelial development and differentiation in luminal A BRCA cells.

**Figure 3:**
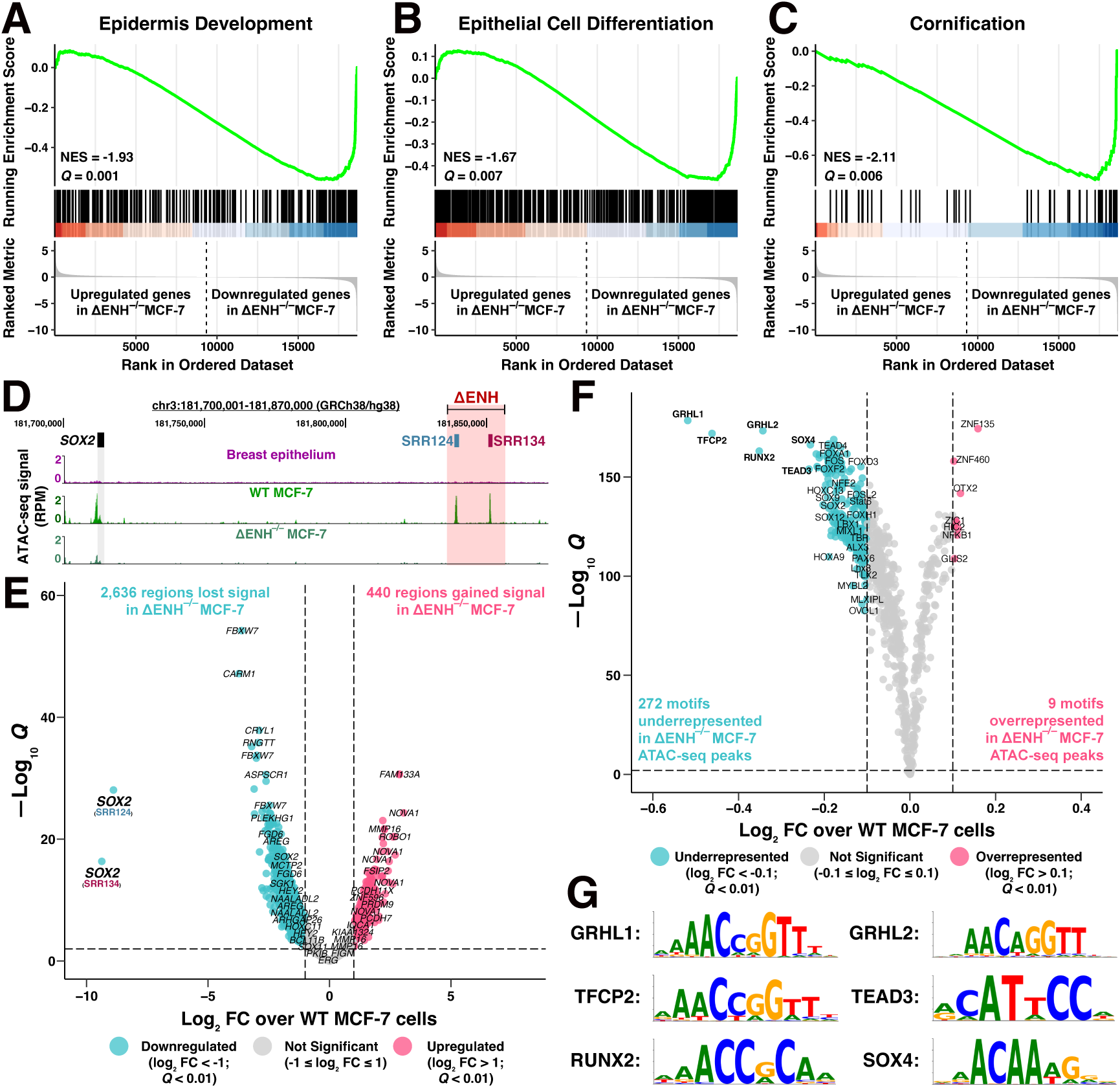
*SOX2* downregulation impacts chromatin accessibility in luminal A BRCA. **(A – C)** Gene Set Enrichment Analysis (GSEA) in the transcriptome of ΔENH^−/−^ compared to WT MCF-7 cells. Genes were ranked according to their change in expression (log_2_ FC). A subset of GO terms significantly enriched among downregulated genes in ΔENH^−/−^ MCF-7 cells are displayed, indicated by the normalized enrichment score (NES) < 1: **(A)** epidermis development, **(B)** epithelial cell differentiation, and **(C)** cornification. GSEA was performed using clusterProfiler (Yu et al. 2012) with an FDR-adjusted *Q* < 0.05 threshold. Green line: running enrichment score. **(D)** UCSC Genome Browser (Kent et al. 2002) view of the SRR124–134 deletion in ΔENH^−/−^ MCF-7 cells with ATAC-seq tracks from breast epithelium (Roadmap Epigenomics Consortium et al. 2015), WT, and ΔENH^−/−^ MCF-7 cells. **(E)** Volcano plot with differential ATAC-seq analysis of ΔENH^−/−^ MCF-7 cells compared to WT. Blue: 2,638 regions that lost (log_2_ FC < -1; FDR-adjusted *Q* < 0.01) chromatin accessibility in ΔENH^−/−^ MCF-7 cells. Pink: 440 regions that gained (log_2_ FC > 1; *Q* < 0.01) chromatin accessibility in ΔENH^−/−^ MCF-7 cells. Grey: 132,726 regions that retained chromatin accessibility in ΔENH^−/−^ MCF-7 cells (-1 ≤ log_2_ FC ≤ 1). Regions were labelled with their closest gene within a ± 1 Mb distance threshold. Differential chromatin accessibility analysis was performed using diffBind (Stark R, Brown G 2011). **(F)** Volcano plot with ATAC-seq footprint analysis of differential transcription factor binding in ΔENH^−/−^ MCF-7 cells compared to WT. Blue: 272 underrepresented (log_2_ FC < -0.1; FDR-adjusted *Q* < 0.01) motifs in ATAC-seq peaks from ΔENH^−/−^ MCF-7 cells. Pink: 9 overrepresented (log_2_ FC > 0.1; *Q* < 0.01) motifs in ATAC-seq peaks from ΔENH^−/−^ MCF-7 cells. Grey: 560 motifs with no representative change (-0.1 ≤ log_2_ FC ≤ 0.1) within ATAC-seq peaks from ΔENH^−/−^ MCF-7 cells. **(G)** Sequence motifs of the top 6 transcription factors with the lowest binding score in ΔENH^−/−^ compared to WT MCF-7 cells: GRHL1, TFCP2, RUNX2, GRHL2, TEAD3, SOX4. Footprint analysis was performed using TOBIAS (Bentsen et al. 2020) utilizing the JASPAR 2022 motif database (Castro-Mondragon et al. 2022).

SOX2 is a pioneer transcription factor that associates with its motif in heterochromatin (Soufi et al. 2012) and recruits chromatin-modifying complexes (Soufi et al. 2015) in embryonic and reprogrammed stem cells. We performed ATAC-seq in ΔENH^−/−^ MCF-7 cells and compared chromatin accessibility to WT MCF-7 cells to identify genome-wide loci that are dependent on SOX2 to remain accessible in luminal A BRCA. As expected, the ATAC-seq signal from all replicates was highly enriched around gene TSS (Supplementary Figure S3A), with both WT and ΔENH^−/−^ samples having higher chromatin accessibility at the TSS of highly expressed genes (Supplementary Figure S3B and Supplementary Figure S3C). Correlation analysis also confirmed the clustering of all three replicates from each genotype (Supplementary Figure S3D). Including the SRR124–134 cluster and pSOX2 (Figure 3D), a total of 3,076 500-bp regions had significant (|log_2_ FC| > 1; FDR-adjusted *Q* < 0.01) changes in chromatin accessibility in ΔENH^−/−^ compared to WT MCF-7 cells (Figure 3E, Supplementary Table S10). Most regions (86%, 2,636 regions) significantly lost chromatin accessibility in ΔENH^−/−^ MCF-7 cells and 76% (2,024 regions) of these regions also gained chromatin accessibility in WT MCF-7 compared to normal breast epithelium (Roadmap Epigenomics Consortium et al. 2015) (Supplementary Table S11). Together, this indicates that SOX2 has an important role in regulating the chromatin accessibility changes acquired in luminal A BRCA.

We used TOBIAS (Bentsen et al. 2020) to analyze changes in transcription factor footprints within ATAC-seq peaks in ΔENH^−/−^ compared to WT MCF-7 cells. From 841 vertebrate motifs (Castro-Mondragon et al. 2022), we found a total of 281 motifs with a significant (|log_2_ FC| > 0.1; FDR-adjusted *Q* < 0.01) differential binding score (Figure 3F, Supplementary Table S12). Most of these motifs (97%, 272 motifs) were underrepresented within ATAC-seq peaks in ΔENH^−/−^ compared to WT MCF-7 cells, indicating that reduced *SOX2* expression affects the binding of multiple other transcription factors. Among them, the GRHL1 (log_2_ FC = -0.519; *Q* = 3×10^-179^), TFCP2 (log_2_ FC = - 0.462; *Q* = 1.03×10^-172^), RUNX2 (log_2_ FC = -0.352; *Q* = 8.02×10^-164^), GRHL2 (log_2_ FC = - 0.343; *Q* = 4.43×10^-174^), TEAD3 (log_2_ FC = -0.235; *Q* = 9.74×10^-155^), and SOX4 (log_2_ FC = -0.232; *Q* = 5.33×10^-167^) motifs (Figure 3G) had the most reduced binding score in ΔENH^−/−^ MCF-7 cells compared to WT MCF-7 cells. These factors belong to three main JASPAR (Castro-Mondragon et al. 2022) motif clusters: GRHL/TFCP (cluster 33; aaAACAGGTTtcAgtt), RUNX (cluster 60; ttctTGtGGTTttt), TEAD (cluster 2; tccAcATTCCAggcCTTta), and SOX (cluster 8; acggaACAATGgaagTGTT). The SOX cluster also included the SOX2 (log_2_ FC = -0.175; *Q* = 6.61×10^-139^) motif.

Next, we aimed to analyze ChIP-seq data from transcription factors within these motif clusters in MCF-7 cells. We utilized two published datasets: GRHL2 (Cocce et al. 2019) and RUNX2 (Jeselsohn et al. 2017). Regions that lost (log_2_ FC < -1; *Q* < 0.01) chromatin accessibility in ΔENH^−/−^ MCF-7 cells significantly (*P* < 2×10^-16^, hypergeometric test) overlapped regions with binding of either of these transcription factors. Among the 2,636 regions that lost chromatin accessibility, 40% (750 regions) also show GRHL2 binding (Supplementary Figure S3E), whereas 21% (552 regions) share RUNX2 binding (Supplementary Figure S3F). In addition, we found multiple SOX motifs significantly (FDR-adjusted *Q* < 0.001) enriched within peaks from both GRHL2 (Supplementary Table S13) and RUNX2 (Supplementary Table S14) ChIP-seq datasets, further suggesting that SOX2 collaborates with GRHL2 and RUNX2 to maintain chromatin accessibility in luminal A BRCA. Expression levels of either *GRHL2* or *RUNX2*, however, were not significantly affected by *SOX2* downregulation in ΔENH^−/−^ MCF-7 cells (-1 ≤ log_2_ FC ≤ 1; Supplementary Table S8), indicating that they are not directly regulated by SOX2 at the transcriptional level but may interact at the protein level.

### The SRR124–134 cluster is associated with *SOX2* overexpression in primary tumors

With the confirmation that the SRR124–134 cluster drives *SOX2* overexpression in the BRCA and LUAD cell lines, we investigated chromatin accessibility at this enhancer cluster within primary tumors isolated from cancer patients. By analyzing the pan-cancer ATAC-seq dataset from TCGA (Corces et al. 2018), we found that SRR124 and SRR134 are most accessible within LUSC, LUAD, BRCA, bladder carcinoma (BLCA), stomach adenocarcinoma (STAD), and uterine endometrial carcinoma (UCEC) patient tumors (Figure 4A). We also quantified the ATAC-seq signal at six other regions (genomic coordinates in Supplementary Table S15): the SOX2 embryonic-associated enhancers (SRR1, SRR2, hSCR), the *SOX2* promoter (pSOX2), a gene regulatory desert with no enhancer features located between the *SOX2* gene and the SRR124–134 cluster (desert), and the promoter of the housekeeping gene *RAB7A* (pRAB7A, positive control). We then compared the chromatin accessibility levels at each of these regions to the promoter of the repressed olfactory gene *OR5K1* (pOR5K1, negative control). Both SRR124 and SRR134 showed significantly increased (*P* < 0.05, Holm-adjusted Dunn’s test) chromatin accessibility when compared to pOR5K1 in BLCA (SRR124 *P* = 0.014; SRR134 *P* = 1.52×10^-3^; Holm-adjusted Dunn’s test), BRCA (SRR124 *P* = 1.70×10^-20^; SRR134 *P* = 1.03×10^-16^), LUAD (SRR124 *P* = 6.76×10^-7^; SRR134 *P* = 3.26×10^-6^), LUSC (SRR124 *P* = 1.62×10^-6^; SRR134 *P* = 7.08×10^-4^), STAD (SRR124 *P* = 1.15×10^-4^; SRR134 *P* = 1.96×10^-7^), and UCEC (SRR124 *P* = 3.15×10^-5^; SRR134 *P* = 0.025) patient tumors (Figure 4B).

**Figure 4:**
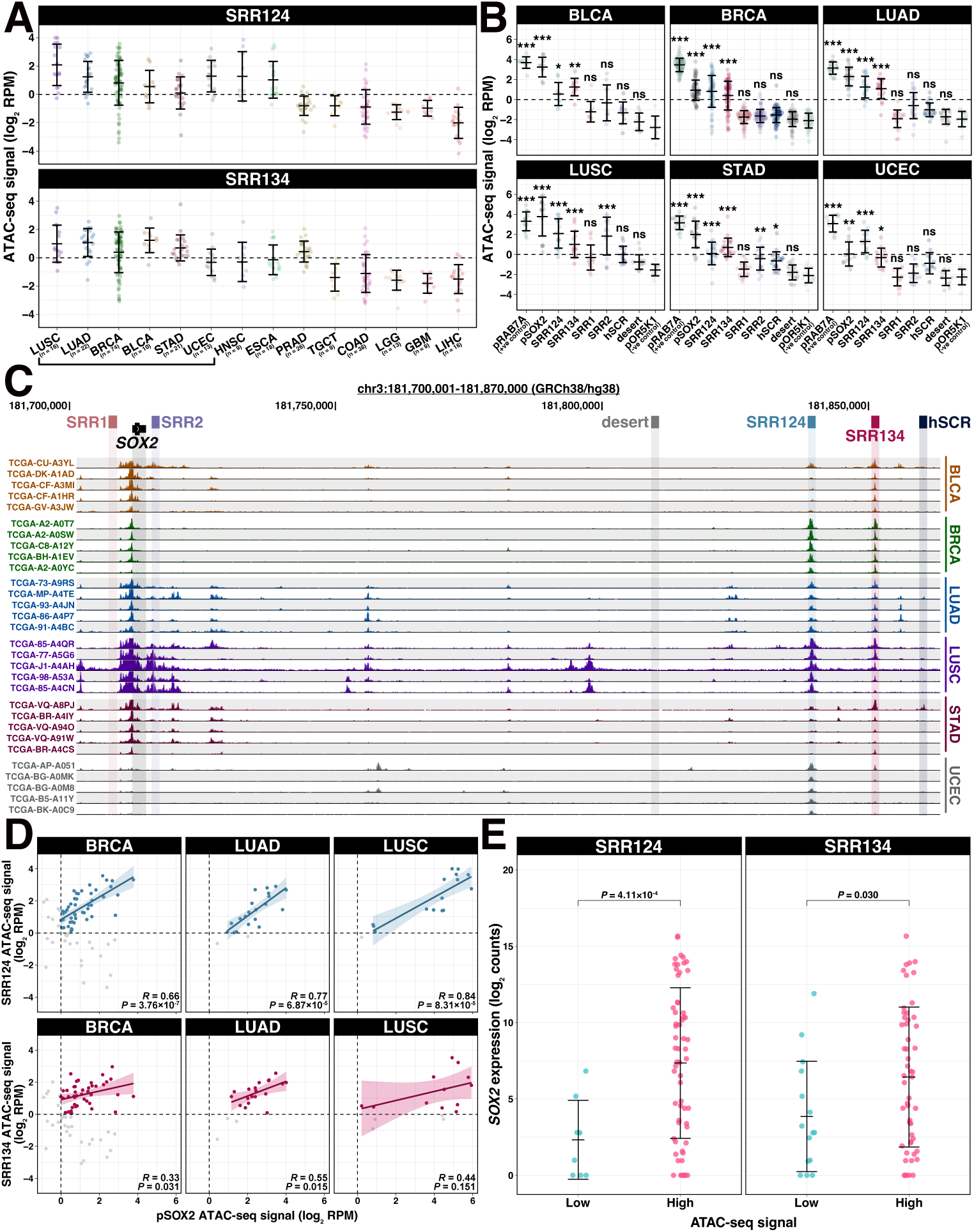
The SRR124–134 cluster is associated with *SOX2* overexpression in cancer patient tumors. **(A)** ATAC-seq signal (log_2_ RPM) at SRR124 and SRR134 for 294 patient tumors from 14 cancer types (Corces et al. 2018). Cancer types are sorted in descending order by the median signal between all three regions. Dashed line: regions with a sum of reads above our threshold (log_2_ RPM > 0) were considered “accessible”. Error bars: standard deviation. Underscore: top 6 cancer types with the highest ATAC-seq median signal. **(B)** ATAC-seq signal (log_2_ RPM) at the *RAB7A* promoter (pRAB7A), *SOX2* promoter (pSOX2), SRR1, SRR2, SRR124, SRR134, hSCR, and a desert region within the *SOX2* locus (desert) compared to the background signal at the repressed *OR5K1* promoter (pOR5K1) in BLCA (n = 10), BRCA (n = 74), LUAD (n = 22), LUSC (n = 16), STAD (n = 21), and UCEC (n = 13) patient tumors. Dashed line: regions with a sum of reads above our threshold (log_2_ RPM > 0) were considered “accessible”. Error bars: standard deviation. Significance analysis by Dunn’s test with Holm correction (* *P* < 0.05, ** *P* < 0.01, *** *P* < 0.001, ns: not significant). **(C)** UCSC Genome Browser (Kent et al. 2002) visualization of the *SOX2* region with ATAC-seq data from BLCA, BRCA, LUAD, LUSC, STAD, and UCEC patient tumors (n = 5 in each cancer type) (Corces et al. 2018). ATAC-seq reads were normalized by library size (RPM). Scale: 0 – 250 RPM. **(D)** ATAC-seq signal at SRR124 and SRR134 regions against ATAC-seq signal for the *SOX2* promoter (pSOX2) from 74 BRCA, 22 LUAD, and 16 LUSC patient tumors. Correlation is shown for accessible chromatin (log_2_ RPM > 0). Grey: tumors with closed chromatin (log_2_ RPM < 0) at either region, not included in the correlation analysis. Significance analysis by Pearson correlation. Bolded line: fitted linear regression model. Shaded area: 95% confidence region for the regression fit. **(E)** Comparison of log_2_-normalized *SOX2* transcript levels (log_2_ counts) between BRCA, LUAD, and LUSC patient tumors according to the chromatin accessibility at SRR124 and SRR134 regions. Chromatin accessibility at each region was considered “low” if log_2_ RPM < -1, or “high” if log_2_ RPM > 1. RNA-seq reads were normalized to library size using DESeq2 (Love et al. 2014). Error bars: standard deviation. Significance analysis by a two-sided t-test with Holm correction.

One explanation for increased chromatin accessibility is locus amplification. While LUSC had high levels of chromatin accessibility likely related to previously described *SOX2* amplifications (Bass et al. 2009; Hussenet et al. 2010; Maier et al. 2011; Liu et al. 2021), most patient tumors showed no evidence of locus amplifications extending to the SRR124–134 cluster, as evidenced by the lack of significant (*P* > 0.05) accessibility at the intermediate desert region. In contrast, the SRR124–134 cluster displayed a consistent pattern of accessible chromatin across multiple cancer types: BLCA, BRCA, LUAD, LUSC, STAD, and UCEC (Figure 4C). GBM and LGG tumors lacked accessible chromatin at this cluster but displayed increased chromatin accessibility at the SRR1 and SRR2 enhancers (Supplementary Figure S4A, Supplementary Table S16), which is consistent with the evidence that SRR1 and SRR2 drive *SOX2* expression in the neural lineage (Ferri et al. 2004; Miyagi et al. 2006; Zappone et al. 2000).

Next, we reasoned that an accessible SRR124–134 cluster drives subsequent *SOX2* transcription within patient tumors. If this is the case, we expect to find positive and significantly correlated chromatin accessibility between this enhancer cluster and pSOX2. Indeed, we found that the majority of BRCA (58%), LUAD (82%), and LUSC (69%) tumors have concurrent accessibility (log_2_ RPM > 0) at pSOX2, SRR124 and SRR134. Patient tumors also showed a significant (*P* < 0.05) correlation (Pearson, *R*) between accessible chromatin signal at pSOX2 and at both SRR124 and SRR134 in BRCA and LUAD (Figure 4D). LUSC tumors showed a significant correlation between accessible chromatin at pSOX2 and SRR124, but not at SRR134 (Figure 4D). As a negative control, we measured the correlation between chromatin accessibility at pSOX2 and at the *SOX2* desert region and found no significant (*P* > 0.05) correlation in any of these cancer types (Supplementary Figure S4B). We also conducted a similar analysis after segregating BRCA tumors into luminal A, luminal B, HER2^+^, and basal-like subtypes (Berger et al. 2018; Corces et al. 2018). Interestingly, we found that both luminal A and luminal B tumors possess a significant (*P* < 0.05) correlation between enhancer accessibility and pSOX2 accessibility, whereas for HER2^+^ tumors the correlation was weaker (Supplementary Figure S4C). Basal-like tumors, on the other hand, display no accessible chromatin at either SRR124 or SRR134. In summary, a luminal-like BRCA phenotype correlates with increased accessibility at the SRR124–134 cluster.

Finally, by separating BRCA, LUAD, and LUSC patient tumors according to their chromatin accessibility at SRR124 and SRR134, we found that tumors with the most accessible chromatin at each of these regions also significantly (*P* < 0.05, t-test) overexpress *SOX2* compared to tumors with low chromatin accessibility at these regions (Figure 4E, Supplementary Table S17). Together, these data are consistent with a model in which increased chromatin accessibility at the SRR124–134 cluster drives *SOX2* overexpression in breast and lung patient tumors.

### *FOXA1* and *NFIB* are upstream regulators of the SRR124–134 cluster

With the indication that the SRR124–134 cluster is driving *SOX2* overexpression in cancer patient tumors, we investigated which transcription factors regulate this cluster in BRCA, LUAD, and LUSC tumors from TCGA (Corces et al. 2018; Hoadley et al. 2014). From a list of 1622 human transcription factors (Lambert et al. 2018), we found that the expression of 115 transcription factors was significantly (FDR-adjusted *Q* < 0.05) associated with chromatin accessibility levels at SRR124, whereas accessibility at SRR134 was associated with the expression of 90 transcription factors (Figure 5A, Supplementary Table S18). From this list, we focused our investigation on FOXA1 and NFIB, which show binding at both SRR124 and SRR134 in ChIP-seq data from MCF-7 cells (ENCODE Project Consortium 2012).

**Figure 5:**
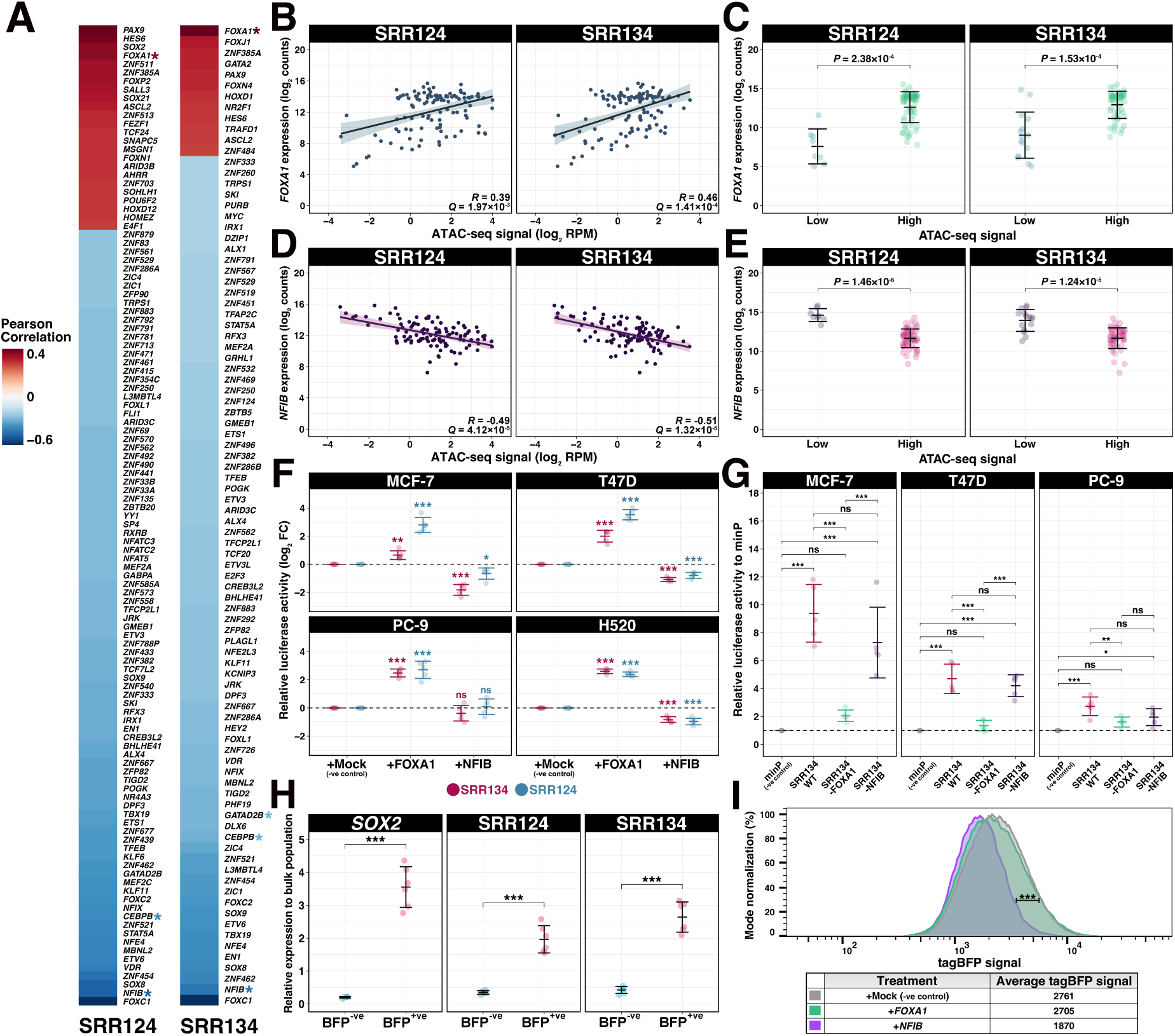
FOXA1 and NFIB are upstream regulators of SRR124 and SRR134. **(A)** Heatmap of the Pearson correlation between transcription factor expression (Hoadley et al. 2014) and chromatin accessibility (Corces et al. 2018) at SRR124 and SRR134 in BRCA, LUAD, and LUSC patient tumors (n = 111). Transcription factors are ordered according to their correlation to chromatin accessibility at each region. Red: transcription factors with a positive correlation (*R* > 0; FDR-adjusted *Q* < 0.05) to chromatin accessibility. Blue: transcription factors with a negative correlation (*R* < 0; *Q* < 0.05) to chromatin accessibility. Asterisk: transcription factors that show binding at SRR124 or SRR134 by ChIP-seq (ENCODE Project Consortium 2012). **(B)** Correlation analysis between *FOXA1* expression (log_2_ counts) and chromatin accessibility (log_2_ RPM) at SRR124 and SRR134 regions in BRCA (n = 74), LUAD (n = 21), and LUSC (n = 16) tumors. RNA-seq reads were normalized to library size using DESeq2 (Love et al. 2014). Significance analysis by Pearson correlation (n = 111). Bolded line: fitted linear regression model. Shaded area: 95% confidence region for the regression fit. **(C)** Comparison of *FOXA1* expression (log_2_ counts) from BRCA, LUAD, and LUSC patient tumors according to their chromatin accessibility at the SRR124 and SRR134 regions. Chromatin accessibility at each region was considered “low” if log_2_ RPM < 1, or “high” if log_2_ RPM > 1. RNA-seq reads were normalized to library size using DESeq2 (Love et al. 2014). Error bars: standard deviation. Significance analysis by a two-sided t-test with Holm correction. **(D)** Correlation analysis between *NFIB* expression (log_2_ counts) and chromatin accessibility (log_2_ RPM) at SRR124 and SRR134 regions in BRCA (n = 74), LUAD (n = 21), and LUSC (n = 16) tumors. RNA-seq reads were normalized to library size using DESeq2 (Love et al. 2014). Significance analysis by Pearson correlation (n = 111). Bolded line: fitted linear regression model. Shaded area: 95% confidence region for the regression fit. **(E)** Comparison of *NFIB* expression (log_2_ counts) from BRCA, LUAD, and LUSC patient tumors according to their chromatin accessibility at the SRR124 and SRR134 regions. Chromatin accessibility at each region was considered “low” if log_2_ RPM < 1, or “high” if log_2_ RPM > 1. RNA-seq reads were normalized to library size using DESeq2 (Love et al. 2014). Error bars: standard deviation. Significance analysis by a two-sided t-test with Holm correction. **(F)** Relative fold change (log_2_ FC) in luciferase activity driven by SRR124 and SRR134 after overexpression of either *FOXA1* or *NFIB* compared to an empty vector containing (mock negative control, miRFP670). Dashed line: average activity of the mock control. Error bars: standard deviation. Significance analysis by Tukey’s test (n = 5; * *P* < 0.05, ** *P* < 0.01, *** *P* < 0.001, ns: not significant). **(G)** Relative luciferase activity driven by WT, FOXA1-mutated, and NFIB-mutated SRR134 constructs compared to a minimal promoter (minP) vector in the MCF-7, PC-9, and T47D cell lines. Dashed line: average activity of minP. Error bars: standard deviation. Significance analysis by Tukey’s test (n = 5; * *P* < 0.05, ** *P* < 0.01, *** *P* < 0.001, ns: not significant). **(H)** RT-qPCR comparison of transcripts at *SOX2*, SRR124, and SRR134 between sorted BFP^-ve^ and BFP^+ve^ MCF-7 cells normalized to unsorted population. Error bars: standard deviation. Significance analysis by paired t-test with Holm correction (n = 6; *** *P* < 0.001). **(I)** FACS density plot comparing tagBFP signal between *SOX2*-P2A-tagBFP MCF-7 cells transfected with an empty vector (mock negative control, miRFP670), FOXA1-T2A-miRFP670, or NFIB-T2A-miRFP670. tagBFP signal was acquired from successfully transfected live cells (miRFP^+^/PI^−^) after 5 days post-transfection. Significance analysis by FlowJo’s chi-squared T(x) test. T(x) scores above 1000 were considered “strongly significant” (*** *P* < 0.001), whereas T(x) scores under 100 were considered “non-significant”.

The expression of *FOXA1* is positively (Pearson correlation *R* > 0) and significantly correlated to accessible chromatin at both SRR124 (*R* = 0.39; FDR-adjusted *Q* = 1.97×10^-3^) and SRR134 (*R* = 0.46; *Q* = 1.41×10^-4^) (Figure 5B). By separating BRCA, LUAD, and LUSC patient tumors according to the chromatin accessibility levels at each region, we found that tumors with the most accessible chromatin within SRR124 (*P* = 2.38×10^-4^, t-test) and SRR134 (*P* = 1.53×10^-4^) also significantly overexpress *FOXA1* compared to tumors with low accessibility at these regions (Figure 5C, Supplementary Table S19). On the other hand, we found the expression of *NFIB* to be negatively (*R* < 0) and significantly correlated with chromatin accessibility at both SRR124 (*R* = -0.49; *Q* = 4.12×10^-5^) and SRR134 (*R* = -0.51; *Q* = 1.32×10^-5^) (Figure 5D). Patient tumors with highly accessible chromatin within SRR124 (*P* = 1.46×10^-6^) and SRR134 (*P* = 1.24×10^-5^) also display significantly downregulated *NFIB* expression (Figure 5E, Supplementary Table S20). These data suggest that whereas FOXA1 could be inducing increased accessibility at the SRR124–134 cluster, NFIB expression could counteract FOXA1 by acting as a repressor. To assess the contribution of these transcription factors to enhancer activity, we overexpressed either *FOXA1* or *NFIB* in H520, MCF-7, PC-9, and T47D cells and compared SRR124 and SRR134 enhancer activity to cells transfected with an empty vector (mock) containing only a fluorescent marker. Although endogenous *FOXA1* and *NFIB* expression levels are already high in both MCF-7 and T47D cells (Supplementary Figure S5A), we found that overexpression of *FOXA1* significantly increased (log_2_ FC > 1; *P* < 0.05, Tukey’s test) the enhancer activity of both SRR124 and SRR134 in the H520, MCF-7, PC-9, and T47D cell lines, whereas *NFIB* overexpression led to a significant decrease (log_2_ FC < 1; *P* < 0.05) in SRR124 and SRR134 enhancer activity in the H520, MCF-7, and T47D cell lines (Figure 5F). This further indicates that *FOXA1* overexpression increases SRR124–134 activity, whereas NFIB represses the enhancer activity of this cluster.

To assess the importance of FOXA1 and NFIB motifs in modulating enhancer activity, we analyzed the SRR134 sequence using the JASPAR2022 motif database (Castro-Mondragon et al. 2022) and mutated FOXA1 (GTAAACA) or NFIB (TGGCAnnnnGCCAA) motifs (mutated SRR134 sequences in Supplementary Table S21). We found that mutation of the FOXA1 motif abolished SRR134 enhancer activity compared to the WT SRR134 sequence within MCF-7 (*P* = 1.53×10^-5^, Tukey’s test), PC-9 (*P* = 1×10^-2^), and T47D (*P* = 4.48×10^-6^) cells, whereas no significant change (*P* > 0.05) in enhancer activity was found for the NFIB-mutated construct (Figure 5G). These data indicate that the FOXA1 motif is crucial for sustaining SRR134 activity, whereas the NFIB motif is dispensable in this context, as would be expected for a negative regulator under conditions where the activity of the target is high.

With the evidence that these transcription factors are modulating SRR124–134 activity, we investigated their transcriptional effects over *SOX2* expression. We used CRISPR homology-directed repair (HDR) to create an MCF-7 cell line in which the *SOX2* gene is tagged with a 2A self-cleaving peptide (P2A) followed by a blue fluorescent protein (tagBFP). This cell line, MCF-7 *SOX2*-P2A-tagBFP, allows rapid visualization of *SOX2* transcriptional changes by measuring tagBFP signal through fluorescence-activated cell sorting (FACS). To validate this model, we sorted cells within the top 10% (BFP^+ve^) and bottom 10% (BFP^-ve^) tagBFP signal (Supplementary Figure S5B). We found that BFP^+ve^ cells showed a significant (*P* = 4.25×10^-5^, paired t-test) increase in *SOX2* expression, and displayed significantly upregulated transcription of enhancer RNA (eRNA) at SRR124 (*P* = 1.54×10^-4^) and SRR134 (*P* = 5.13×10^-5^) compared to BFP^-ve^ cells (Figure 5H). This confirms that the tagBFP signal is directly correlated to *SOX2* transcription levels and enhancer output in MCF-7 *SOX2*-P2A-tagBFP cells.

Finally, we overexpressed *FOXA1* or *NFIB* in MCF-7 SOX2-P2A-tagBFP to assess changes in *SOX2* transcription. Although overexpression of *FOXA1* did not significantly (chi-squared T(x)=63.70) change tagBFP signal, we found that overexpression of *NFIB* significantly (chi-squared T(x)=1168.88) reduced tagBFP signal compared to transfection of an empty vector (mock) (Figure 5I). This confirms the repressive effect of NFIB over *SOX2* expression and illustrates a potential mechanism upstream of *SOX2* that modulates chromatin accessibility at the SRR124–134 cluster and subsequent control of *SOX2* transcription in cancer cells.

### SRR124 and SRR134 are conserved enhancers across mammals and are required for the separation of the anterior foregut

*SOX2* is required for the proper development of multiple tissues (Arnold et al. 2011), including the digestive and respiratory systems in the mouse (Francis et al. 2019; Gontan et al. 2008; Que et al. 2007, 2009; Teramoto et al. 2020; Tompkins et al. 2009) and in humans (Zenteno et al. 2006; Williamson et al. 2006; Chassaing et al. 2007). Therefore, we questioned whether the SRR124–134 cluster drives *SOX2* expression in additional contexts other than cancer. A compilation of chromatin accessibility data from cardiac, digestive, embryonic, lymphoid, musculoskeletal, myeloid, neural, placental, pulmonary, renal, skin, and vascular tissues (ENCODE Project Consortium 2012; Meuleman et al. 2020; Roadmap Epigenomics Consortium et al. 2015) showed that both SRR124 and SRR134 display increased chromatin accessibility in digestive and respiratory tissues alongside cancer samples (Figure 6A). By comparing DNase-seq signal from fetal lung and stomach tissues (ENCODE Project Consortium 2012), we found that both SRR124 (lung *P* = 1.25×10^-6^; stomach *P* = 9.64×10^-4^; Holm-adjusted Dunn’s test) and SRR134 (lung *P* = 1.14×10^-3^; stomach *P* = 0.045), together with SRR2 (lung *P* = 1.55×10^-3^; stomach *P* = 5.74×10^-5^), are significantly more accessible than pOR5K1 (Figure 6B, Supplementary Table S22). This suggests that SRR124 and SRR134 are contributing to *SOX2* expression during the development of the digestive and respiratory systems.

**Figure 6:**
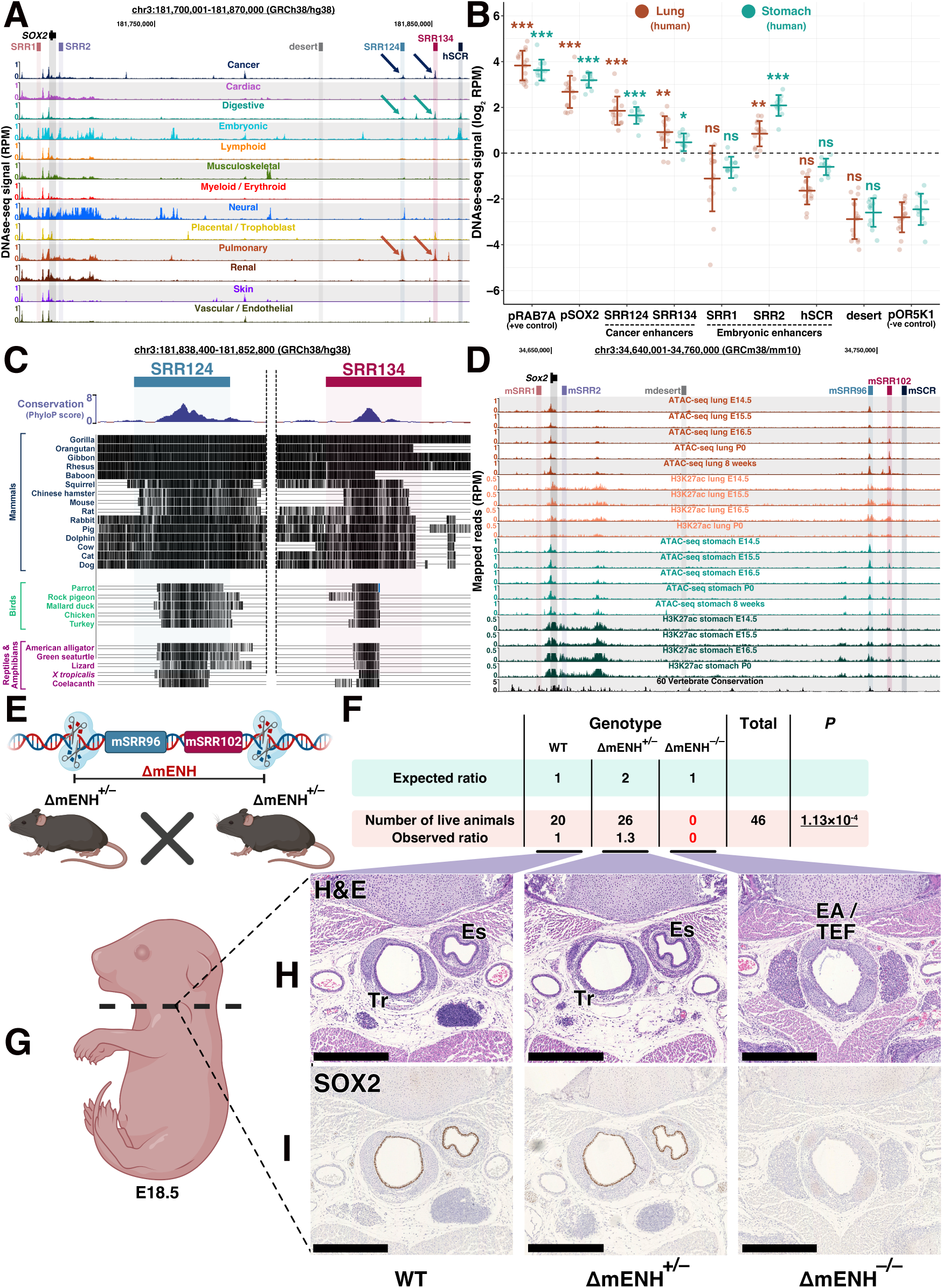
The SRR124 and SRR134 enhancers are conserved across species and are required for the separation of the esophagus and trachea in the mouse. **(A)** UCSC Genome Browser (Kent et al. 2002) view of the *SOX2* region containing a compilation of chromatin accessibility tracks of multiple human tissues (ENCODE Project Consortium 2012; Meuleman et al. 2020; Roadmap Epigenomics Consortium et al. 2015). Arrow: increased chromatin accessibility at the SRR124–134 cluster in cancer, digestive, and respiratory tissues. **(B)** DNAse-seq quantification (log_2_ RPM) at the *RAB7A* promoter (pRAB7A), *SOX2* promoter (pSOX2), SRR1, SRR2, SRR124, SRR134, human SCR (hSCR), and a desert region within the *SOX2* locus (desert) compared to the background signal at the repressed *OR5K1* promoter (pOR5K1) in lung and stomach embryonic tissues (ENCODE Project Consortium 2012). Dashed line: Regions with a sum of reads above our threshold (log_2_ RPM > 0) were considered “accessible”. Error bars: standard deviation. Significance analysis by Dunn’s test with Holm correction (* *P* < 0.05, ** *P* < 0.01, *** *P* < 0.001, ns: not significant). **(C)** UCSC Genome Browser (Kent et al. 2002) with PhyloP conservation scores (Pollard et al. 2010) at the SRR124 and SRR134 enhancers across mammals, birds, reptiles, and amphibians species. Black lines: highly conserved sequences. Empty lines: variant sequences. **(D)** UCSC Genome Browser (Kent et al. 2002) view of the *Sox2* region in the mouse. ATAC-seq and H3K27ac ChIP-seq data from lung and stomach tissues throughout developmental days E14.5 to the 8^th^ post-natal week (ENCODE Project Consortium 2012; Liu et al. 2019). mSRR96: homologous to SRR124. mSRR102: homologous to SRR134. Reads were normalized to library size (RPM). **(E)** Illustration demonstrating the mSRR96–102 enhancer cluster CRISPR deletion (ΔmENH) in C57BL/6J mouse embryos. **(F)** Quantification and genotype of the C57BL/6J progeny from mSRR96–102-deleted crossings (ΔmENH^+/–^). Pups were counted and genotyped at weaning (P21). Significance analysis by chi-squared test to measure the deviation in the number of obtained pups from the expected mendelian ratio of 1:2:1 (WT : ΔmENH^+/–^ : ΔmENH^−/−^). **(G)** Transverse cross-section of fixed E18.5 embryos at the start of the thymus. **(H)** Embryo sections stained with Hematoxylin and Eosin (H&E). Scale bar: 500µm. Es: esophagus; Tr: trachea; EA/TEF: esophageal atresia with distal tracheoesophageal fistula. **(I)** Embryo cross-sections stained for SOX2. Scale bar: 500µm. Es: esophagus; Tr: trachea; EA/TEF: esophageal atresia with distal tracheoesophageal fistula.

Since critical developmental genes are often controlled by highly conserved enhancers across species (Pennacchio et al. 2006; Woolfe et al. 2005), we hypothesized that the SRR124–134 cluster might regulate *SOX2* expression during the development of other species. By analyzing PhyloP conservation scores (Pollard et al. 2010; Kent et al. 2002), we discovered that both SRR124 and SRR134 contain a highly conserved core sequence that is preserved across mammals, birds, reptiles, and amphibians (Figure 6C). After aligning and comparing enhancer sequences between humans and mice, we found the core sequence at both SRR124 and SRR134 are highly conserved (> 80%) in the mouse genome (Supplementary Figure S6A). We termed these homologous regions as mSRR96 (96 kb downstream of the mouse *Sox2* promoter; homologous to the human SRR124) and mSRR102 (102 kb downstream of the mouse *Sox2* promoter; homologous to the human SRR134). Enhancer feature analysis in the developing lung and stomach tissues in the mouse (ENCODE Project Consortium 2012; Liu et al. 2019) showed that both mSRR96 and mSRR102 display increased chromatin accessibility and H3K27ac signal throughout developmental days E14.5 to the 8^th^ post-natal week (Figure 6D). Interestingly, mSRR96 and mSRR102 display higher ATAC-seq and H3K27ac signal towards the later stages of development in the lungs, but at early stages of development in the stomach. This suggests a distinct spatiotemporal contribution of this homologous cluster to *Sox2* expression during the development of these tissues in the mouse. ATAC-seq quantification (genomic coordinates in Supplementary Table S23) showed that both mSRR96 (lung *P* = 5.54×10^-5^; stomach *P* = 2.37×10^-4^; Holm-adjusted Dunn’s test) and mSRR102 (lung *P* = 1.27×10^-3^; stomach *P* = 0.046) are significantly more accessible than the repressed promoter of the olfactory gene *Olfr266* (pOlfr266, negative control) during the development of the lungs and stomach in the mouse (Supplementary Figure S6B, Supplementary Table S24). Together, these results suggest a conserved *SOX2* regulatory mechanism across multiple species and support a model in which the SRR124 and SRR134 enhancers and their homologs regulate *SOX2* expression during the development of the digestive and respiratory systems.

To assess the contribution of the mSRR96 and mSRR102 regions to the development of the mouse, we generated a knockout containing a deletion spanning the mSRR96–102 enhancer cluster (ΔmENH) (Figure 6E). We crossed animals carrying a heterozygous mSRR96–102 deletion (ΔmENH^+/–^) and determined the number of pups alive at weaning (P21) from each genotype. We found a significant (*P* = 1.13×10^-4^, Chi-squared test) deviation from the expected mendelian ratio with no homozygous mice (ΔmENH^−/−^) alive at weaning (Figure 6F), demonstrating that the mSRR96–102 enhancer cluster is crucial for survival in the mouse. To investigate the resulting phenotype in a homozygous mSRR96–102 enhancer deletion, we collected E18.5 embryos and prepared cross-sections at the thymus level from five animals of each genotype (WT, ΔmENH^+/–^, and ΔmENH^−/−^) (Figure 6G). Similar to other studies that interfered with *Sox2* expression during development (Que et al. 2007; Teramoto et al. 2020; Chakraborty et al. 2023), we found that all five ΔmENH^−/−^ embryos developed EA/TEF, where the esophagus and trachea fail to separate during embryonic development (Figure 6H). WT and ΔmENH^+/–^ embryos, on the other hand, showed normal development of the esophageal and tracheal tissues. Immunohistochemistry showed the complete absence of the SOX2 protein within the EA/TEF tissue in ΔmENH^−/−^ embryos, whereas WT and ΔmENH^+/–^ embryos showed high levels of SOX2 protein within both the esophagus and tracheal tubes (Figure 6I). Finally, immunofluorescence staining for NKX2.1, a transcription factor associated with the inner epithelium of the respiratory tract (Minoo et al. 1999), showed high protein levels within the inner layer of the EA/TEF tissue in ΔmENH^−/−^ embryos, indicating that this aberrant tissue resembles a tracheal-like structure lacking SOX2 (Supplementary Figure S6C). Together, these results demonstrate that mSRR96 and mSRR102 are required to drive *Sox2* expression during the development and separation of the esophagus and trachea.

## DISCUSSION

Our findings reveal that the SRR124–134 enhancer cluster is essential for *Sox2* expression in the developing digestive and respiratory systems as it is required for the separation of the esophagus and trachea during mouse development. When embryogenesis is complete, *Sox2* expression is downregulated in most differentiated cell types as its developmental enhancers are decommissioned. We propose that aberrant upregulation of the pioneer factor *FOXA1* recommissions both SRR124 and SRR134 in tumor cells, driving *SOX2* overexpression in breast and lung cancer. As SOX2 is also a pioneer transcription factor, increased levels of this protein further reprogram the chromatin landscape of cancer cells, binding at multiple downstream regulatory regions, increasing chromatin accessibility, and driving subsequent upregulation of genes associated with epithelium development, ultimately supporting a tumor-initiating phenotype.

The observation that enhancers involved in the development of the digestive and respiratory systems are reprogrammed to support *SOX2* upregulation during tumorigenesis is in line with previous observations that tumor-initiating cells acquire a less differentiated phenotype (Bonnet and Dick 1997; Chaffer et al. 2011; Lapidot et al. 1994, 199; Gupta et al. 2011). It is more surprising, however, that the *SOX2* gene is regulated by common enhancers in both breast and lung cancer cells as enhancers are usually highly tissue-specific (Pennacchio et al. 2006; Woolfe et al. 2005; Thurman et al. 2012; Stergachis et al. 2013). Our observation that *FOXA1* expression is significantly correlated to chromatin accessibility at the SRR124–134 cluster and increases the transcriptional output of the SRR124 and SRR134 enhancers provides a mechanistic link between breast and lung developmental programs and cancer progression. FOXA1 is directly involved in the branching morphogenesis of the epithelium in breast (Bernardo et al. 2010; Liu et al. 2016) and lung (Besnard et al. 2005, 1; Paranjapye et al. 2020) tissues, where SOX2 also plays an important role (Domenici et al. 2019; Gontan et al. 2008). Overexpression of both *FOXA1* (Camolotto et al. 2018; Fu et al. 2016, 2019; Lupien et al. 2008; Orstad et al. 2022; Roe et al. 2017; Stergachis et al. 2013) and *SOX2* (Boumahdi et al. 2014; Chou et al. 2013, 2; Liu et al. 2013) have also been individually linked to the activation of transcriptional programs associated with multiple types of cancer. Therefore, we propose that FOXA1 is one of the key players responsible for the reprogramming of the SRR124–134 cluster in cancer, which then drives *SOX2* overexpression in breast and lung tumors. As mutation of the FOXA1 motif disrupted SRR134 enhancer activity, and this motif is shared among other members of the forkhead box (FOX) transcription factor family (Pierrou et al. 1994), it remains unclear if FOXA1 alone activates the SRR124–134 cluster, or whether other FOX proteins are involved in this process. For example, *FOXM1* overexpression, which also showed binding at both SRR124 and SRR134 in MCF-7 cells, has similarly been associated with poor patient outcomes in multiple types of cancer (Li et al. 2017).

In addition to the activating role of FOXA1, we identified NFIB as a negative regulator of *SOX2* expression through inhibition of SRR124–134 activity. NFIB is normally required for the development of multiple tissues (reviewed in Harris et al. 2015), including the brain and lungs (Steele-Perkins et al. 2005; Gründer et al. 2002; Hsu et al. 2011), tissues in which *SOX2* expression is also tightly regulated (Favaro et al. 2009; Gontan et al. 2008). In the lungs, NFIB is essential for promoting the maturation and differentiation of progenitor cells (Gründer et al. 2002; Steele-Perkins et al. 2005). This is in stark contrast to SOX2, which inhibits the differentiation of lung cells (Gontan et al. 2008). Interestingly, NFIB seems to have paradoxical roles in cancer, acting both as a tumor suppressor and as an oncogene in different tissues (Becker-Santos et al. 2017). Among its tumor suppressor activity, NFIB acts as a barrier to skin carcinoma progression (Zhou et al. 2014b), and its downregulation is associated with dedifferentiation and aggressiveness in LUAD (Becker-Santos et al. 2016). On the other hand, SOX2 promotes skin (Boumahdi et al. 2014) and lung (Ferone et al. 2016) cancer progression. As an oncogene, *NFIB* promotes cell proliferation and metastasis in STAD (Wu et al. 2018), where *SOX2* downregulation is associated with poor patient outcomes (Otsubo et al. 2008; Wang et al. 2015; Zhang et al. 2010). With this contrasting relationship between *SOX2* and *NFIB* across multiple tissues, we propose that NFIB normally acts as a suppressor of SRR124–134 activity and *SOX2* expression during the differentiation of progenitor cells; downregulation of *NFIB* expression then results in *SOX2* overexpression during tumorigenesis of the breast and lung.

We initially hypothesized that SRR1 and SRR2 (Miyagi et al. 2004; Zappone et al. 2000; Tomioka et al. 2002), and/or the SCR (Li et al. 2014; Zhou et al. 2014a) might be recommissioned during cancer progression, as stem cell-related enhancers have been shown to acquire enhancer features in tumorigenic cells (Aran et al. 2016). Although other studies have also proposed the activation of either SRR1 (Leis et al. 2012; Stolzenburg et al. 2012) or SRR2 (Iglesias et al. 2014; Jung et al. 2015) as the main drivers of *SOX2* overexpression in BRCA, we found no evidence of this mechanism and instead identified the SRR124–134 cluster as the main driver of *SOX2* expression in BRCA and LUAD. Our patient tumor analysis did show that GBM and LGG were the only cancer types that display a unique and consistent pattern of accessible chromatin at SRR1 and SRR2, which is likely related to glioma cells assuming a neural stem cell-like identity to sustain high levels of cell proliferation in the brain (Bulstrode et al. 2017). In fact, SRR2 deletion was shown to downregulate *SOX2* and reduce cell proliferation in GBM cells (Saenz-Antoñanzas et al. 2021), highlighting enhancer specificity to different tumor types. In line with these findings, our observation that PC-9 LUAD cells are dependent on SRR124– 134 for *SOX2* transcription, whereas in H520 LUSC cells SRR124–134 is dispensable, again highlights these tumor-type specific regulatory mechanisms. LUSC tumors frequently amplify the *SOX2* locus (Bass et al. 2009; Hussenet et al. 2010; Liu et al. 2021; Maier et al. 2011), whereas LUAD tumors do not (Björkqvist et al. 1998), indicating that different mechanisms are involved in genome dysregulation in these two subtypes of lung cancer. Interestingly, a further downstream enhancer cluster located ∼55 kb away from SRR124–134 is co-amplified with *SOX2* in LUSC cell lines (Liu et al. 2021), revealing an additional mechanism that could sustain *SOX2* overexpression in the absence of the SRR124–134 cluster in certain types of LUSC but not in LUAD.

Deletion of mSRR96–102, homolog of the human SRR124–134 cluster, resulted in EA/TEF which is also observed in human cases with *SOX2* heterozygous mutations (Zenteno et al. 2006; Williamson et al. 2006; Chassaing et al. 2007). Interestingly, a recent study showed that insertion of a CTCF insulation cluster downstream of the *Sox2* gene, but upstream of mSRR96–102, disrupts *Sox2* expression, impairs separation of the esophagus and trachea, and results in perinatal lethality due to EA/TEF in the mouse (Chakraborty et al. 2023). This was of particular interest for understanding enhancer functional nuances during development since the SCR, which is required for *Sox2* transcription at implantation, can partially overcome the insulator effect of this insertion. The authors proposed that enhancer density might explain the EA/TEF phenotype, as chromatin features suggested that enhancers in the developing lung and stomach tissues might be spread over a 400 kb domain (Chakraborty et al. 2023). The 6 kb deletion that removes the mSRR96–102 cluster causing EA/TEF suggests this is not the case. Instead, we propose that the sensitivity of each cell type to gene dosage is behind the differing ability of CTCF to block distal enhancers. This is based on two observations: in humans, heterozygous *SOX2* mutations are linked with the anophthalmia-esophageal-genital syndrome (Zenteno et al. 2006; Williamson et al. 2006; Chassaing et al. 2007); in mice, hypomorphic *Sox2* alleles display similar phenotypes in the eye (Taranova et al. 2006) and EA/TEF (Que et al. 2007; Teramoto et al. 2020). This suggests that cells from the peri-implantation phase are less sensitive to lower *Sox2* dosages compared to cells from the developing airways and digestive systems in both species and explains the aberrant phenotypes observed at term.

Overall, our findings illustrate how cis-regulatory regions can similarly drive gene expression in both normal and diseased contexts and serve as a prime example of how decommissioned developmental enhancers may be reprogrammed during tumorigenesis. The fact that we have found a digestive/respiratory-associated enhancer cluster driving gene expression in a non-native context such as BRCA remains intriguing and reinforces a model in which tumorigenic cells often revert to a progenitor-like state that combines cis-regulatory features of progenitor cells from multiple developing lineages (Stergachis et al. 2013). This “dys-differentiation” mechanism seems to be centered around the overexpression of a few key development-associated pioneer transcription factors such as FOXA1 and SOX2. Identifying additional mechanisms that regulate the reprogramming of these enhancers could lead to new approaches to target tumor-initiating cells that depend on *SOX2* overexpression.

## MATERIALS AND METHODS

### Cell Culture

MCF-7 cells were obtained from Eldad Zacksenhaus (Toronto General Hospital Research Institute, Toronto, CA). H520 (HTB-182) and T47D (HTB-133) cells were acquired from ATCC. PC-9 (90071810) cells were obtained from Sigma. Cell line identities were confirmed by short tandem repeat profiling. MCF-7 and T47D cells were grown in phenol red-free DMEM high glucose (Gibco), 10% FBS (Gibco), 1x Glutamax (Gibco), 1x Sodium Pyruvate (Gibco), 1x Penicillin-Streptomycin (Gibco), 1x Non-essential amino acids (Gibco), 25 mM HEPES (Gibco) and 0.01 mg/ml insulin (Sigma). H520 and PC-9 cells were grown in phenol red-free RPMI-1640 (Gibco), 10% FBS (Gibco), 1x Glutamax (Gibco), 1x Sodium Pyruvate (Gibco), 1x Penicillin-Streptomycin (Gibco), 1x Non-essential amino acids (Gibco), and 25 mM HEPES (Gibco). Cells were either passaged or had their medium replenished every three days.

### Genome editing

Pairs of gRNA plasmids were constructed by inserting a 20 bp target sequence (Supplementary Table S25) into an empty gRNA cloning vector (a gift from George Church; Addgene plasmid # 41824; http://n2t.net/addgene:41824; RRID:Addgene_41824) (Mali et al. 2013, 9) containing either miRFP670 (Addgene plasmid #163748) or tagBFP (Addgene plasmid #163747) fluorescent markers. Plasmids were sequenced to confirm correct insertion. Both gRNA (1 µg each) vectors were co-transfected with 3 µg of pCas9_GFP (a gift from Kiran Musunuru; Addgene plasmid #44719; http://n2t.net/addgene:44719; RRID:Addgene_44719) (Ding et al. 2013) using Neon electroporation (Life Technologies). After 72 hours of transfection, cells were FACS sorted to select clones that contained all three plasmids. Sorted tagBFP^+^/GFP^+^/miRFP670^+^ cells were grown in a bulk population and serially diluted into individual wells to generate isogenic populations. Once fully grown, each well was screened by PCR to confirm the deletion.

### Gene tagging

*SOX2* was tagged with a P2A-tagBFP sequence in both alleles using CRISPR-mediated homology-directed repair (HDR) (Ran et al. 2013). This strategy results in the expression of a single transcript that is further translated into two separate proteins due to ribosomal skipping (Ahier and Jarriault 2014). In summary, we designed a gRNA that targets the 3’ end of the *SOX2* stop codon (Supplementary Table S25, Addgene plasmid #163752). We then amplified ∼800 bp homology arms upstream and downstream of the gRNA target sequence using high-fidelity Phusion Polymerase. We purposely avoided amplification of the *SOX2* promoter sequence to reduce the likelihood of random integrations in the genome. Both homology arms were then joined at each end of a P2A-tagBFP sequence using Gibson assembly. Flanking primers containing the gRNA target sequence were used to reamplify *SOX2*-P2A-tagBFP and add gRNA targets at both ends of the fragment; this approach allows excision of the HDR sequence from the backbone plasmid once inside the cell (Zhang et al. 2017). Finally, the full HDR sequence was inserted into a pJET1.2 (Thermo Scientific) backbone, midiprepped, and sequenced (Addgene #163751). 3µg of HDR template was then co-transfected with 1µg of hCas9 (a gift from George Church; Addgene plasmid #41815; http://n2t.net/addgene:41815 ; RRID:Addgene_41815) (Mali et al. 2013) and 1µg of gRNA plasmid using Neon electroporation (Life Technologies). A week after transfection, tagBFP^+^ cells were FACS sorted as a bulk population. Sorted cells were further grown for two more weeks, and single tagBFP^+^ cells were isolated to generate isogenic populations. Once fully grown, each clone was screened by PCR and sequenced to confirm homozygous integration of P2A-tagBFP into the *SOX2* locus.

### Luciferase assay

Luciferase activity was measured using the dual-luciferase reporter assay (Promega #E1960) that relies on the co-transfection of two plasmids: pGL4.23 (Firefly Luciferase, *luc2*) and pGL4.75 (Renilla Luciferase). Assayed plasmids were constructed by subcloning the empty pGL4.23 vector containing a minimal promoter (minP). SRR124, SRR134, SRR1, SRR2, and hSCR were PCR-amplified (primers in Supplementary Table S26) from MCF-7 genomic DNA using high-fidelity Phusion Polymerase and inserted in the forward position downstream of the *luc2* gene at the NotI restriction site. Constructs were sequenced to confirm correct insertions.

JASPAR2022 (Castro-Mondragon et al. 2022) was used to find FOXA1 (GTAAACA) and NFIB (TGGCAnnnnGCCAA) motifs in the SRR134 sequence. Only motifs with a score of 80% or higher were further analyzed. Bases within each motif sequence were mutated until the score was reduced below 80% without affecting co-occurring motifs or creating novel binding sites. In total, four FOXA1 motifs and two NFIB motifs were mutated (Supplementary Table S21). Engineered sequences were ordered as gene blocks (Eurofins) and inserted into pGL4.23 in the forward position. Constructs were sequenced to confirm correct insertions.

Cells were plated in 96-well plates with 4 technical replicates at 2.10^4^ cells per well. After 24 hours, a 200ng 50:1 mixture of enhancer vector and pGL4.75 was transfected using Lipofectamine 3000 (0.05µl Lipofectamine:1µl Opti-mem). For transcription factor overexpression analysis, a 200ng 50:10:1 mixture of enhancer vector, expression plasmid, and pGL4.75 was transfected. After 48 hours of transfection, cells were lysed in 1x Passive Lysis Buffer and stored at -80°C until all 5 biological replicates were completed. Luciferase activity was measured in the Fluoroskan Ascent FL plate reader. Enhancer activity was calculated by normalizing the firefly signal from pGL4.23 to the *Renilla* signal from pGL4.75.

### Colony formation assay

MCF-7 and PC-9 cells were seeded at low density (2,000 cells/well) into 6-well plates in triplicate for each cell line. Culture media was renewed every 3 days. After 12 days, cells were fixed with 3.7% paraformaldehyde for 10 minutes and stained with 0.5% crystal violet for 20 minutes to quantify the number of colonies formed. Crystal violet staining was then eluted with 10% acetic acid and absorbance was measured at 570 nm to evaluate cell proliferation. Each 6-well plate was considered one biological replicate and the experiment was repeated five times for each cell line (n = 5).

### FACS analysis

For analyzing the effects of *FOXA1* and *NFIB* overexpression, 2.10^6^ SOX2-P2A-tagBFP cells were transfected with 50nM of plasmid expressing either miRFP670 (a gift from Vladislav Verkhusha; Addgene plasmid #79987; http://n2t.net/addgene:79987; RRID:Addgene_79987), FOXA1-T2A-miRFP670 (Addgene plasmid #182335), or NFIB-T2A-miRFP670 (Addgene plasmid #187222) in 5 replicates. Five days after transfection, miRFP670, tagBFP, and propidium iodide (PI) (live/dead stain) signals were acquired; the amount of tagBFP signal from miRFP670^+^/PI^−^ cells was compared between each treatment across all replicates.

FlowJo’s chi-squared T(x) test was used to contrast the effects of each treatment over tagBFP expression; T(x) scores above 1000 were considered “strongly significant” (***), whereas T(x) scores under 100 were considered “non-significant”.

### Transcriptome analysis

Total RNA was isolated from WT and enhancer-deleted (ΔENH) cell lines using the RNeasy kit. Genomic DNA was digested by Turbo DNAse. 500-2,000ng of total RNA was used in a reverse transcription reaction with random primers. cDNA was diluted in H_2_O and amplified in a qPCR reaction using SYBR Select Mix (primers in Supplementary Table S27). Amplicons were sequenced to confirm primer specificity. Gene expression was normalized to *PUM1* (Kılıç et al. 2014; Krasnov et al. 2019; Lyng et al. 2008).

Total RNA was sent to The Centre for Applied Genomics (TCAG) for paired-end rRNA-depleted total RNA-seq (Illumina 2500, 125 bp). Read quality was checked by fastQC, trimmed using fastP (Chen et al. 2018) and mapped to the human genome (GRCh38/hg38) using STAR 2.7 (Dobin et al. 2013). Normal breast epithelium RNA-seq was obtained from ENCODE (Supplementary Table S28) (ENCODE Project Consortium 2012; Roadmap Epigenomics Consortium et al. 2015). Mapped reads were quantified using featureCounts (Liao et al. 2014) and imported into DESeq2 (Love et al. 2014) for normalization and differential expression analysis. Genes with a |log_2_ FC| > 1 and FDR-adjusted *Q* < 0.01 were considered significantly changing. Differential gene expression was plotted using the EnhancedVolcano package. Correlation and clustering heatmaps were plotted using the pheatmap R package (https://cran.r-project.org/web/packages/pheatmap/index.html). Signal enrichment plot was prepared using NGS.plot (Shen et al. 2014).

Cancer patient transcriptome data were obtained from TCGA (Hoadley et al. 2014) using the TCGAbiolinks package (Colaprico et al. 2016). The overall survival KM-plot was calculated using clinical information from TCGA (Liu et al. 2018a). Tumor transcriptome data were compared to normal tissue using DESeq2. RNA-seq reads were normalized to library size using DESeq2 (Love et al. 2014) and transformed to a log_2_ scale [log_2_ counts]. Differential gene expression was considered significant if |log_2_ FC| > 1 and *Q* < 0.01.

Gene Set Enrichment Analysis (GSEA) was performed by ranking genes according to their log_2_ FC in ΔENH^−/−^ versus WT MCF-7 cells. The ranking was then analyzed using the GSEA function from the clusterProfiler package (Yu et al. 2012) with a threshold of FDR-adjusted *Q* < 0.05 using the MSigDB GO term database (C5).

### Chromatin accessibility analysis

Cells were grown in three separate wells (n = 3) and 50,000 cells were sent to Princess Margaret Genomics Centre for ATAC-seq library preparation using the Omni-ATAC protocol (Corces et al. 2017). ATAC-seq libraries were sequenced using 50 bp paired-ended parameters in the Illumina Novaseq 6000 platform. Read quality was checked by fastQC, trimmed using fastP and mapped to the human genome (GRCh38/hg38) using STAR 2.7. narrowPeaks were called using Genrich (https://github.com/jsh58/Genrich). Differential chromatin accessibility analysis was performed using diffBind (Stark R, Brown G 2011). ATAC-seq peaks with a |log_2_ FC| > 1 and FDR-adjusted *Q* < 0.01 were considered significantly changing. Correlation heatmaps were generated using diffBind. Signal enrichment plot was prepared using NGS.plot (Shen et al. 2014). Genes were separated into three categories according to their expression levels in our WT MCF-7 RNA-seq data.

Transcription factor footprint analysis was performed using TOBIAS (Bentsen et al. 2020) with standard settings. Motifs with a |log_2_ FC| > 0.1 and FDR-adjusted *Q* < 0.01 were considered significantly enriched in each condition. Replicates (n = 3) were merged into a single BAM file for each treatment. Motif enrichment at differential ATAC-seq peaks was performed using HOMER (Heinz et al. 2010). ATAC-seq peaks were assigned to their closest gene within ± 1 Mb distance from their promoter using ChIPpeakAnno (Zhu et al. 2010).

Cancer patient ATAC-seq data was obtained from TCGA (Corces et al. 2018). DNAse-seq from human developing tissues were obtained from ENCODE (Supplementary Table S28) (ENCODE Project Consortium 2012; Roadmap Epigenomics Consortium et al. 2015). Read quantification was calculated at the *RAB7a* (pRAB7a), *OR5K1* (pOR5K1), and *SOX2* (pSOX2) promoters, together with SRR1, SRR2, SRR124, SRR134, hSCR, and desert regions with a 1,500 bp window centered at the core of each region (genomic coordinates of each region in Supplementary Table S15). Reads were normalized to library size (RPM) and transformed to a log_2_ scale (log_2_ RPM) using a custom script (https://github.com/luisabatti/BAMquantify). Each region’s average log_2_ RPM was compared to the *OR5K1* promoter for differential analysis using Dunn’s test with Holm correction. Correlations were calculated using Pearson’s correlation test and considered significant if FDR-adjusted *Q* < 0.05. Chromatin accessibility at SRR124 and SRR134 regions was considered low if log_2_ RPM < -1, medium if -1 ≤ log_2_ RPM ≤ 1, or high if log_2_ RPM > 1.

ATAC-seq from developing mouse lung and stomach tissues were obtained from ENCODE (Supplementary Table S28) (ENCODE Project Consortium 2012) and others (Liu et al. 2019). Conserved mouse regulatory regions were lifted from the human build (GRCh38/hg38) to the mouse build (GRCm38/mm10) using UCSC liftOver (Kent et al. 2002, 200). The number of mapped reads was calculated at the *Egf* (pEgf), *Olfr266* (pOlfr266), and *Sox2* (pSox2) promoters, together with the mouse mSRR1, mSRR2, mSRR96, mSRR102, mSCR and desert regions with a 1,500 bp window at each location (genomic coordinates in Supplementary Table S23). Each log2-transformed region’s reads per million (log_2_ RPM) was compared to the negative *Olfr266* promoter control for differential analysis using Dunn’s test with Holm correction.

### Conservation analysis

Cross-species evolutionary conservation was obtained using phyloP (Pollard et al. 2010). Pairwise comparisons between human SRR124 and SRR134 (GRCh38/hg38) and mouse mSRR96 and mSRR102 (GRCm38/mm10) sequences were plotted using FlexiDot (Seibt et al. 2018) with an 80% conservation threshold.

### ChIP-seq analysis

Transcription factor and histone modifications ChIP-seq were obtained from ENCODE (ENCODE Project Consortium 2012) (Supplementary Table S28) and others (Chan et al. 2018; Cocce et al. 2019; Sato et al. 2019) (Supplementary Table S29). H3K4me1 and H3K27ac tracks were normalized to input and library size (log_2_ RPM). ATAC-seq reads were normalized to library size (RPM). Histone modification ChIP-seq tracks and transcription factor ChIP-seq peaks were uploaded to the UCSC browser (Kent et al. 2002, 20) for visualization. Normalized H3K4me1, H3K27ac and ATAC-seq reads were quantified and the difference in normalized signal was calculated using diffBind. Peaks with a |log_2_ FC| > 1 and *Q* < 0.01 were considered significantly changing.

Overlapping ChIP-seq and ATAC-seq peaks were analyzed using ChIPpeakAnno (Zhu et al. 2010). The hypergeometric test was performed by comparing the number of overlapping peaks to the total size of the genome divided by the median peak size.

### Mouse line construction

Our mSRR96–102 knockout mouse line (C57BL/6J-Del(3)1Tcp/Jmit) was ordered from and generated by The Centre for Phenogenomics (TCP) in Toronto, ON. The protocol for the generation of the mouse line has been previously described (Gertsenstein and Nutter 2021). Briefly, C57BL/6J zygotes were collected from superovulated, mated, and plugged female mice at 0.5-day post coitum. Zygotes were electroporated with Cas9 RNPs complexes (gRNA sequences in Supplementary Table S25) and transferred into pseudopregnant female recipients within 3-4 hours of electroporation. Born pups (founders) were screened by end-point PCR and sequenced to confirm allelic mSRR96– 102 deletions. Heterozygous mSRR96–102 founders (ΔmENH^+/–^) were then backcrossed to the parental strain to confirm germline transmission. Once the mouse line was established and the mSRR96–102 deletion was fully confirmed and sequenced in the N1 offspring, ΔmENH^+/–^ mice were crossed and the number of live pups from each genotype (WT, ΔmENH^+/–^, ΔmENH^−/−^) was assessed at weaning (P21). The obtained number of live pups from each genotype was then compared to the expected mendelian ratio of 1:2:1 (WT : ΔmENH^+/–^ : ΔmENH^−/−^) using a chi-squared test. Once the lethality of the homozygous deletion was confirmed at weaning, E18.5 embryos generated from new ΔmENH^+/–^ crosses were collected for further histological analyses. All procedures involving animals were performed in compliance with the Animals for Research Act of Ontario and the Guidelines of the Canadian Council on Animal Care. The TCP Animal Care Committee reviewed and approved all procedures conducted on animals at the facility.

### Histological analyses

A total of 46 embryos were collected at E18.5 and fixed in 4% paraformaldehyde. Each embryo was genotyped, and a total of 15 embryos, 5 of each genotype (WT, ΔmENH^+/–^, ΔmENH^−/−^), were randomly selected, processed, and embedded in paraffin for sectioning and further analysis. Tissue sections were collected at 4µm thickness roughly at the start of the thymus. Sections were prepared by the Pathology Core at TCP.

Tissue sections were stained with Hematoxylin and Eosin (H&E) using an auto-stainer to ensure batch consistency. Slides were scanned using a Hamamatsu Nanozoomer slide scanner at 20X magnification. For immunohistochemistry staining, E18.5 embryo cross-sections were submitted to heat-induced epitope retrieval with TRIS-EDTA (pH 9.0) for 10 minutes, followed by quenching of endogenous peroxidase with Bloxall reagent (Vector). Non-specific antibody binding was blocked with 2.5 % normal horse serum (Vector), followed by incubation for 1 hour in Rabbit anti-SOX2 (Abcam, ab92494, 1:500). After washes, sections were incubated for 30 minutes with ImmPRESS Anti-Rabbit HRP (Vector), followed by DAB reagent and counterstained in Mayer’s hematoxylin.

For immunofluorescence staining, E18.5 embryo cross-sections were collected onto charged slides and then baked at 60 °C for 30 minutes. Tissue sections were submitted to heat-induced epitope retrieval with Citrate Buffer pH 6.0 for 10 minutes. Non-specific antibody binding was blocked with Protein Block Serum-Free (Dako) for 10 minutes, followed by overnight incubation at 4 °C in a primary antibody cocktail (Rabbit anti-NKX2.1, Abcam ab76013 at 1:200; Rat anti-SOX2, Thermo Fisher Scientific 14-9811-80 at 1:100). After washes with TBS-T, sections were incubated for 1 hour with a cocktail of Alexa Fluor-conjugated secondary antibodies at 1:200 (Goat anti-Rabbit IgG AF488 – Thermo Fisher Scientific A32731; Goat anti-Rat IgG AF647 – Thermo Fisher Scientific A21247), followed by counterstaining with DAPI. Scanning was performed using an Olympus VS-120 slide scanner and imaged using a Hamamatsu ORCA-R2 C10600 digital camera for all dark-field and fluorescent images.

## SUPPORTING INFORMATION

**Supplementary Figure S1: (A)** Super-logarithmic volcano plot of *PUM1* expression from RNA-seq of 21 cancer types compared to normal tissue (Hoadley et al. 2014). Cancer types with log_2_ FC > 1 and FDR-adjusted *Q* < 0.01 were considered to significantly overexpress *PUM1*. Error bars: standard deviation. **(B)** Kaplan-Meier plot (Kaplan and Meier 1958) of overall survival against time since diagnosis for 3,064 patients with BRCA (n = 1089), COAD (n = 453), GBM (n = 153), LIHC (n = 370), LUAD (n = 504), and LUSC (n = 495) tumors (Liu et al. 2018a). We divided patients into four equal groups and compared two groups: high *SOX2* expression (range: 10.06–16.36 log_2_ counts) and low *SOX2* expression (range: 0–1.67 log_2_ counts). RNA-seq reads were normalized to library size using DESeq2 (Love et al. 2014). Significance analysis by logrank test. The shadowed area represents the 95% confidence interval. **(C)** Comparison of *SOX2* expression (log_2_ counts) between luminal A (n = 560), luminal B (n = 207), HER2+ (n = 82), basal-like (n = 190) breast cancer subtypes (Berger et al. 2018; Hoadley et al. 2014) and normal mammary tissue (n = 152) (Hoadley et al. 2014). RNA-seq reads were normalized to library size using DESeq2 (Love et al. 2014). Error bars: standard deviation. Significance analysis by Tukey’s test (*** *P* < 0.001, * *P* < 0.05, ns: not significant). **(D)** Volcano plot with DESeq2 (Love et al. 2014, 2) differential expression analysis between WT MCF-7 cells and breast epithelium (Roadmap Epigenomics Consortium et al. 2015). Blue: 7,937 genes that significantly lost expression (log_2_ FC < -1; FDR-adjusted *Q* < 0.01) in WT MCF-7 cells. Pink: 5,335 genes that significantly gained expression (log_2_ FC > 1; *Q* < 0.01) in WT MCF-7 cells. Grey: 25,342 genes that maintained similar (-1 ≤ log_2_ FC ≤ 1) expression between WT MCF-7 and breast epithelium cells. **(E)** RT-qPCR analysis of *SOX2* transcript levels in the MCF-7, T47D, PC-9 and H520 cell lines. Error bars: standard deviation. Significance analysis by Tukey’s test (n = 3; *** *P* < 0.001).

**Supplementary Figure S2: (A)** SOX2 protein levels in mouse embryonic stem cells (mESC, positive control), WT, ΔENH^+/–^, and ΔENH^−/−^ MCF-7 clones. Cyclophilin A (CypA) was used as a loading control across all samples. **(B)** RT-qPCR analysis of *SOX2* transcript levels in SRR124–134 heterozygous- (ΔENH^+/–^) and homozygous- (ΔENH^−/−^) deleted H520 (LUSC) clones compared to WT cells. Error bars: standard deviation. Significance analysis by Dunnett’s test (n = 3, ns: not significant). **(C)** Euclidean distance pairwise comparison between WT and ΔENH^−/−^ MCF-7 replicates (n = 3) using variance stabilizing transformed RNA-seq reads from DESeq2 (Love et al. 2014). Darker colors indicate a higher correlation. **(D)** Euclidean hierarchical clustering of 529 differentially expressed genes (|log_2_ FC| > 1; FDR-adjusted *Q* < 0.01) based on RNA-seq analysis between WT and ΔENH^−/−^ MCF-7 replicates (n = 3). Reads were normalized for each gene across treatments (Z-score). Blue color indicates downregulated genes (Z-score < 0). Red color indicates upregulated genes (Z-score > 0).

**Supplementary Figure S3: (A)** ATAC-seq metagene enrichment plot ± 2 kb around the transcription start site (TSS) across all genes from WT and ΔENH^−/−^ MCF-7 cells (n = 3). Reads were normalized by library size (RPM). Grey: TSS. Shaded area: standard deviation. **(B)** ATAC-seq metagene enrichment plot ± 2 kb around the transcription start site (TSS) across 12,167 high (average log_2_ counts = 9.12), 12,167 medium (average log_2_ counts = 2.94), and 12,167 low (average log_2_ counts = 0.43) expressed genes in WT MCF-7 cells. Genes were split into each group according to RNA-seq data. RNA-seq reads were normalized to library size using DESeq2 (Love et al. 2014). Grey: TSS. (**C)** ATAC-seq metagene enrichment plot ± 2 kb around the transcription start site (TSS) across 12,167 high (average log_2_ counts = 9.10), 12,167 medium (average log_2_ counts = 2.97), and 12,167 low (average log_2_ counts = 0.47) expressed genes in ΔENH^−/−^ MCF-7 cells. Genes were split into each group according to RNA-seq data. RNA-seq reads were normalized to library size using DESeq2 (Love et al. 2014). Grey: TSS. **(D)** Pairwise Pearson correlation comparison between WT and ΔENH^−/−^ MCF-7 replicates (n = 3) using ATAC-seq normalized signal from diffBind (Stark R, Brown G 2011). Darker colors indicate a higher correlation. **(E)** Overlap between GRHL2 ChIP-seq peaks (Cocce et al. 2019) and ATAC-seq peaks that significantly (log_2_ FC < -1; *P* < 0.01) lost chromatin accessibility in ΔENH^−/−^ MCF-7 cells. Significance analysis by the hypergeometric test (Zhu et al. 2010). **(F)** Overlap between RUNX2 ChIP-seq peaks (Jeselsohn et al. 2017) and ATAC-seq peaks that significantly (log_2_ FC < -1; *P* < 0.01) lost chromatin accessibility in ΔENH^−/−^ MCF-7 cells. Significance analysis by the hypergeometric test (Zhu et al. 2010).

**Supplementary Figure S4: (A)** ATAC-seq signal at the *RAB7A* promoter (pRAB7A), *SOX2* promoter (pSOX2), SRR1, SRR2, SRR124, SRR134, human SCR (hSCR) and desert region versus the background signal at the repressed *OR5K1* promoter (pOR5K1) in BLCA (n = 10), BRCA (n = 74), COAD (n = 38), ESCA (n = 18), GBM (n = 9), HNSC (n = 9), LGG (n = 13), LIHC (n = 16), LUAD (n = 22), LUSC (n = 16), PRAD (n = 26), STAD (n = 21), TGCT (n = 9), and UCEC (n = 13) patient tumors. Dashed line: regions with log_2_ RPM > 0 were considered “accessible”. Error bars: standard deviation. Significance analysis by Dunn’s test with Holm correction (* *P* < 0.05, ** *P* < 0.01, *** *P* < 0.001, ns: not significant). **(B)** ATAC-seq signal at the *SOX2* desert region (desert) against ATAC-seq signal for the *SOX2* promoter (pSOX2) from 74 BRCA, 22 LUAD, and 16 LUSC patient tumors. Dashed line: regions with log_2_ RPM > 0 were considered “accessible”. Significance analysis by Pearson correlation. Bolded line: fitted linear regression model. Shaded area: 95% confidence region for the regression fit. **(C)** ATAC-seq signal at SRR124 and SRR134 regions against ATAC-seq signal for the *SOX2* promoter (pSOX2) from BRCA patient tumors separated into luminal A (n = 31), luminal B (n = 16), HER2^+^ (n = 10), and basal-like (n = 14) subtypes (Berger et al. 2018; Corces et al. 2018). Correlation is shown for accessible chromatin (log_2_ RPM > 0). Grey: tumors with closed chromatin (log_2_ RPM < 0) at either region, not included in the correlation analysis. Significance analysis by Pearson correlation. Bolded line: fitted linear regression model. Shaded area: 95% confidence region for the regression fit.

**Supplementary Figure S5: (A)** RT-qPCR analysis of *FOXA1 and NFIB* expression in the H520, MCF-7, PC-9, and T47D cell lines. Error bars: standard deviation. Significance analysis by Tukey’s test (n = 3; ** *P* < 0.01, *** *P* < 0.001). **(B)** FACS plot of tagBFP signal (450 nm) over side scatter (SSC) in WT and *SOX2*-P2A-tagBFP MCF-7 cells. Cell populations within the top 10% tagBFP signal were considered “tagBFP positive” (BFP^+ve^), whereas populations within the bottom 10% BFP signal were considered “tagBFP negative” (BFP^-ve^).

**Supplementary Figure S6: (A)** Dot-plot alignment of human (GRCh38/hg38, y-axis) SRR124 and SRR134, and mouse (GRCm38/mm10, x-axis) mSRR96 and mSRR102 homologous sequences (1,500 bp). Lines indicate high conservation scores (> 80%) across both species. Sequence alignment using Clustal Omega (Sievers et al. 2011). **(B)** ATAC-seq quantification (log_2_ RPM) at the promoter of the housekeeping gene *Egf* (pEgf, positive control), *Sox2* promoter (pSox2), mSRR1, mSRR2, mSRR96, mSRR102, mSCR, and a mouse desert (mdesert) region compared to the background signal at the repressed *Olfr266* promoter (pOlfr266) in lung and stomach embryonic tissues from the mouse (ENCODE Project Consortium 2012). mSRR96: homologous to SRR124. mSRR102: homologous to SRR134. Dashed line: regions with a sum of reads above our threshold (log_2_ RPM > 0) were considered “accessible”. Error bars: standard deviation. Significance analysis by Dunn’s test with Holm correction (* *P* < 0.05, ** *P* < 0.01, *** *P* < 0.001, ns: not significant). **(C)** Immunofluorescence staining of WT, ΔENH^+/–^, and ΔENH^−/−^ E18.5 embryos with DAPI (blue), NKX2.1 (green), and SOX2 (white). Cross-sections were prepared at the thymus levels. Scale bar: 200µm. Es: esophagus; Tr: trachea; EA/TEF: esophageal atresia with distal tracheoesophageal fistula.

**Supplementary Table S1:** List of TCGA tumor type abbreviations.

**Supplementary Table S2:** *SOX2* differential expression analysis between primary tumor vs. normal tissue across TCGA cancer types.

**Supplementary Table S3:** *PUM1* differential expression analysis between primary tumor vs. normal tissue across TCGA cancer types.

**Supplementary Table S4:** TCGA cancer patient overall survival analysis relative to *SOX2* expression levels.

**Supplementary Table S5:** TCGA copy number variation (CNV) and *SOX2* expression analysis.

**Supplementary Table S6:** RNA-seq differential expression analysis between WT MCF-7 vs. breast epithelium (ENCODE).

**Supplementary Table S7:** Differential ATAC-seq, H3K4me1, and H3K27ac analysis within ± 1 Mb of the *SOX2* gene in WT MCF-7 vs. Breast epithelium (ENCODE).

**Supplementary Table S8:** RNA-seq differential expression analysis comparing ΔENH^−/−^ versus WT MCF-7 cells.

**Supplementary Table S9:** Gene set enrichment analysis (GSEA) in WT versus ΔENH^−/−^ MCF-7 cells.

**Supplementary Table S10:** Significantly changing ATAC-seq peaks in ΔENH^−/−^ versus WT MCF-7 cells.

**Supplementary Table S11:** ATAC-seq peaks that commonly gained signal in WT MCF-7 vs. breast epithelium and lost signal in ΔENH^−/−^ MCF-7 cells.

**Supplementary Table S12:** ATAC-seq footprint analysis in ΔENH^−/−^ vs. WT MCF-7 cells.

**Supplementary Table S13:** ChIP-seq motif analysis of GRHL2 peaks in WT MCF-7 cells.

**Supplementary Table S14:** ChIP-seq motif analysis of RUNX2 peaks in WT MCF-7 cells.

**Supplementary Table S15:** Coordinates of regions used in genome-wide analysis in humans (GRCh38/hg38).

**Supplementary Table S16:** Chromatin accessibility analysis across TCGA cancer types.

**Supplementary Table S17:** ATAC-seq quantification used to separate patient tumors into expression groups and their *SOX2* expression levels.

**Supplementary Table S18:** Significantly correlated transcription factors to accessible chromatin at the SRR124–134 cluster in BRCA, LUAD, and LUSC tumors.

**Supplementary Table S19:** *FOXA1* transcript levels and chromatin accessibility at the SRR124–134 cluster in BRCA, LUAD, and LUSC patient tumors.

**Supplementary Table S20:** *NFIB* transcript levels and chromatin accessibility at the SRR124–134 cluster in BRCA, LUAD, and LUSC patient tumors.

**Supplementary Table S21:** WT, FOXA1, and NFIB mutated SRR134 sequences.

**Supplementary Table S22:** Chromatin accessibility analysis in human (GRCh38/hg38) lung and stomach embryonic tissues.

**Supplementary Table S23:** Coordinates of regions used in genome-wide analysis in the mouse (GRCm38/mm10).

**Supplementary Table S24:** Chromatin accessibility analysis in mouse (GRCm38/mm10) embryonic lung and stomach tissues

**Supplementary Table S25:** List of gRNA sequences used for CRISPR/Cas9.

**Supplementary Table S26:** List of primers used for enhancer cloning.

**Supplementary Table S27:** List of primers used in RT-qPCR experiments.

**Supplementary Table S28:** ENCODE datasets used in this paper.

**Supplementary Table S29:** GEO datasets used in this paper.

## DATA AVAILABILITY

Sequencing and processed data files were submitted to the Gene Expression Omnibus (GEO; https://www.ncbi.nlm.nih.gov/geo/) repository (GSE132344).

## ACKNOWLEDGMENTS

We thank all the members of the Mitchell laboratory for helpful discussions and Mathieu Lupien for manuscript review. We also thank the ENCODE Consortium and the TCGA project for generating and releasing data to the scientific community. Finally, we thank the contribution of the staff at TCP (The Centre for Phenogenomics), including Kyle Roberton, who handled the embedding, cutting, and H&E staining of E18.5 mouse embryos, and Vivian Bradaschia, responsible for the IHC staining. This work was supported by the Canadian Institutes of Health Research (FRN PJT153186 and PJT180312), the Canada Foundation for Innovation, and the Ontario Ministry of Research and Innovation (operating and infrastructure grants held by J.A.M.). BioRender.com was used to create parts of Figures 6E and 6G.

## AUTHOR CONTRIBUTIONS

L.E.A. designed and performed bioinformatic analyses, cell culture work, CRISPR deletions, data curation, gene expression quantification, and led the conceptualization and writing of the manuscript; P.L.F. assessed cellular phenotypes, including the colony formation assay; L.H. acquired and processed TCGA ATAC-seq data, and assisted in writing review & editing; M.C. assisted in the writing review & editing; M.M.H. provided TCGA data access and assisted in writing review & editing; J.A.M. was involved in supervision, funding acquisition, data interpretation, experimental design, and writing review & editing. All authors have participated in the editing and approval of the manuscript.

## REFERENCES

Ahier A, Jarriault S. 2014. Simultaneous Expression of Multiple Proteins Under a Single Promoter in *Caenorhabditis elegans* via a Versatile 2A-Based Toolkit. Genetics 196: 605–613.

Alonso MM, Diez-Valle R, Manterola L, Rubio A, Liu D, Cortes-Santiago N, Urquiza L, Jauregi P, de Munain AL, Sampron N, et al. 2011. Genetic and Epigenetic Modifications of Sox2 Contribute to the Invasive Phenotype of Malignant Gliomas ed. B.W. Futscher. PLoS ONE 6: e26740.

Aran D, Abu-Remaileh M, Levy R, Meron N, Toperoff G, Edrei Y, Bergman Y, Hellman A. 2016. Embryonic Stem Cell (ES)-Specific Enhancers Specify the Expression Potential of ES Genes in Cancer. PLoS Genet 12: e1005840.

Arnold K, Sarkar A, Yram MA, Polo JM, Bronson R, Sengupta S, Seandel M, Geijsen N, Hochedlinger K. 2011. Sox2(+) adult stem and progenitor cells are important for tissue regeneration and survival of mice. Cell Stem Cell 9: 317–329.

Avilion AA. 2003. Multipotent cell lineages in early mouse development depend on SOX2 function. Genes & Development 17: 126–140.

Bass AJ, Watanabe H, Mermel CH, Yu S, Perner S, Verhaak RG, Kim SY, Wardwell L, Tamayo P, Gat-Viks I, et al. 2009. SOX2 is an amplified lineage-survival oncogene in lung and esophageal squamous cell carcinomas. Nature Genetics 41: 1238– 1242.

Becker-Santos DD, Lonergan KM, Gronostajski RM, Lam WL. 2017. Nuclear Factor I/B: A Master Regulator of Cell Differentiation with Paradoxical Roles in Cancer. EBioMedicine 22: 2–9.

Becker-Santos DD, Thu KL, English JC, Pikor LA, Martinez VD, Zhang M, Vucic EA, Luk MT, Carraro A, Korbelik J, et al. 2016. Developmental transcription factor NFIB is a putative target of oncofetal miRNAs and is associated with tumour aggressiveness in lung adenocarcinoma. J Pathol 240: 161–172.

Bentsen M, Goymann P, Schultheis H, Klee K, Petrova A, Wiegandt R, Fust A, Preussner J, Kuenne C, Braun T, et al. 2020. ATAC-seq footprinting unravels kinetics of transcription factor binding during zygotic genome activation. Nat Commun 11: 4267.

Berezovsky AD, Poisson LM, Cherba D, Webb CP, Transou AD, Lemke NW, Hong X, Hasselbach LA, Irtenkauf SM, Mikkelsen T, et al. 2014. Sox2 promotes malignancy in glioblastoma by regulating plasticity and astrocytic differentiation. Neoplasia 16: 193–206, 206.e19–25.

Berger AC, Korkut A, Kanchi RS, Hegde AM, Lenoir W, Liu W, Liu Y, Fan H, Shen H, Ravikumar V, et al. 2018. A Comprehensive Pan-Cancer Molecular Study of Gynecologic and Breast Cancers. Cancer Cell 33: 690–705.e9.

Bernardo GM, Lozada KL, Miedler JD, Harburg G, Hewitt SC, Mosley JD, Godwin AK, Korach KS, Visvader JE, Kaestner KH, et al. 2010. FOXA1 is an essential determinant of ERalpha expression and mammary ductal morphogenesis. Development 137: 2045–2054.

Besnard V, Wert SE, Kaestner KH, Whitsett JA. 2005. Stage-specific regulation of respiratory epithelial cell differentiation by Foxa1. Am J Physiol Lung Cell Mol Physiol 289: L750–759.

Bi M, Zhang Z, Jiang Y-Z, Xue P, Wang H, Lai Z, Fu X, De Angelis C, Gong Y, Gao Z, et al. 2020. Enhancer reprogramming driven by high-order assemblies of transcription factors promotes phenotypic plasticity and breast cancer endocrine resistance. Nat Cell Biol 22: 701–715.

Björkqvist AM, Husgafvel-Pursiainen K, Anttila S, Karjalainen A, Tammilehto L, Mattson K, Vainio H, Knuutila S. 1998. DNA gains in 3q occur frequently in squamous cell carcinoma of the lung, but not in adenocarcinoma. Genes Chromosomes Cancer 22: 79–82.

Bonnet D, Dick JE. 1997. Human acute myeloid leukemia is organized as a hierarchy that originates from a primitive hematopoietic cell. Nat Med 3: 730–737.

Boumahdi S, Driessens G, Lapouge G, Rorive S, Nassar D, Le Mercier M, Delatte B, Caauwe A, Lenglez S, Nkusi E, et al. 2014. SOX2 controls tumour initiation and cancer stem-cell functions in squamous-cell carcinoma. Nature 511: 246–250.

Boyle AP, Davis S, Shulha HP, Meltzer P, Margulies EH, Weng Z, Furey TS, Crawford GE. 2008. High-Resolution Mapping and Characterization of Open Chromatin across the Genome. Cell 132: 311–322.

Brunner HG, van Bokhoven H. 2005. Genetic players in esophageal atresia and tracheoesophageal fistula. Curr Opin Genet Dev 15: 341–347.

Buenrostro JD, Giresi PG, Zaba LC, Chang HY, Greenleaf WJ. 2013. Transposition of native chromatin for multimodal regulatory analysis and personal epigenomics. Nat Methods 10: 1213–1218.

Bulstrode H, Johnstone E, Marques-Torrejon MA, Ferguson KM, Bressan RB, Blin C, Grant V, Gogolok S, Gangoso E, Gagrica S, et al. 2017. Elevated FOXG1 and SOX2 in glioblastoma enforces neural stem cell identity through transcriptional control of cell cycle and epigenetic regulators. Genes Dev 31: 757–773.

Camolotto SA, Pattabiraman S, Mosbruger TL, Jones A, Belova VK, Orstad G, Streiff M, Salmond L, Stubben C, Kaestner KH, et al. 2018. FoxA1 and FoxA2 drive gastric differentiation and suppress squamous identity in NKX2-1-negative lung cancer. Elife 7: e38579.

Carter D, Chakalova L, Osborne CS, Dai Y, Fraser P. 2002. Long-range chromatin regulatory interactions in vivo. Nature Genetics 32: 623.

Castro-Mondragon JA, Riudavets-Puig R, Rauluseviciute I, Berhanu Lemma R, Turchi L, Blanc-Mathieu R, Lucas J, Boddie P, Khan A, Manosalva Pérez N, et al. 2022. JASPAR 2022: the 9th release of the open-access database of transcription factor binding profiles. Nucleic Acids Research 50: D165–D173.

Chaffer CL, Brueckmann I, Scheel C, Kaestli AJ, Wiggins PA, Rodrigues LO, Brooks M, Reinhardt F, Su Y, Polyak K, et al. 2011. Normal and neoplastic nonstem cells can spontaneously convert to a stem-like state. PNAS 108: 7950–7955.

Chakraborty S, Kopitchinski N, Zuo Z, Eraso A, Awasthi P, Chari R, Mitra A, Tobias IC, Moorthy SD, Dale RK, et al. 2023. Enhancer–promoter interactions can bypass CTCF-mediated boundaries and contribute to phenotypic robustness. Nat Genet. https://www.nature.com/articles/s41588-022-01295-6 (Accessed February 1, 2023).

Chan HL, Beckedorff F, Zhang Y, Garcia-Huidobro J, Jiang H, Colaprico A, Bilbao D, Figueroa ME, LaCava J, Shiekhattar R, et al. 2018. Polycomb complexes associate with enhancers and promote oncogenic transcriptional programs in cancer through multiple mechanisms. Nat Commun 9: 3377.

Chassaing N, Gilbert-Dussardier B, Nicot F, Fermeaux V, Encha-Razavi F, Fiorenza M, Toutain A, Calvas P. 2007. Germinal mosaicism and familial recurrence of a SOX2 mutation with highly variable phenotypic expression extending from AEG syndrome to absence of ocular involvement. Am J Med Genet A 143A: 289–291.

Chen C, Morris Q, Mitchell JA. 2012. Enhancer identification in mouse embryonic stem cells using integrative modeling of chromatin and genomic features. BMC Genomics 13: 152.

Chen S, Zhou Y, Chen Y, Gu J. 2018. fastp: an ultra-fast all-in-one FASTQ preprocessor. Bioinformatics 34: i884–i890.

Chen Y, Shi L, Zhang L, Li R, Liang J, Yu W, Sun L, Yang X, Wang Y, Zhang Y, et al. 2008. The Molecular Mechanism Governing the Oncogenic Potential of SOX2 in Breast Cancer. Journal of Biological Chemistry 283: 17969–17978.

Chew J-L, Loh Y-H, Zhang W, Chen X, Tam W-L, Yeap L-S, Li P, Ang Y-S, Lim B, Robson P, et al. 2005. Reciprocal Transcriptional Regulation of Pou5f1 and Sox2 via the Oct4/Sox2 Complex in Embryonic Stem Cells. Mol Cell Biol 25: 6031–6046.

Chou Y-T, Lee C-C, Hsiao S-H, Lin S-E, Lin S-C, Chung C-H, Chung C-H, Kao Y-R, Wang Y-H, Chen C-T, et al. 2013. The Emerging Role of SOX2 in Cell Proliferation and Survival and Its Crosstalk with Oncogenic Signaling in Lung Cancer. STEM CELLS 31: 2607–2619.

Cocce KJ, Jasper JS, Desautels TK, Everett L, Wardell S, Westerling T, Baldi R, Wright TM, Tavares K, Yllanes A, et al. 2019. The Lineage Determining Factor GRHL2 Collaborates with FOXA1 to Establish a Targetable Pathway in Endocrine Therapy-Resistant Breast Cancer. Cell Rep 29: 889–903.e10.

Colaprico A, Silva TC, Olsen C, Garofano L, Cava C, Garolini D, Sabedot TS, Malta TM, Pagnotta SM, Castiglioni I, et al. 2016. TCGAbiolinks: an R/Bioconductor package for integrative analysis of TCGA data. Nucleic Acids Research 44: e71–e71.

Corces MR, Granja JM, Shams S, Louie BH, Seoane JA, Zhou W, Silva TC, Groeneveld C, Wong CK, Cho SW, et al. 2018. The chromatin accessibility landscape of primary human cancers. Science 362: eaav1898.

Corces MR, Trevino AE, Hamilton EG, Greenside PG, Sinnott-Armstrong NA, Vesuna S, Satpathy AT, Rubin AJ, Montine KS, Wu B, et al. 2017. An improved ATAC-seq protocol reduces background and enables interrogation of frozen tissues. Nature Methods 14: 959–962.

Cox JL, Wilder PJ, Desler M, Rizzino A. 2012. Elevating SOX2 levels deleteriously affects the growth of medulloblastoma and glioblastoma cells. PLoS One 7: e44087.

Creyghton MP, Cheng AW, Welstead GG, Kooistra T, Carey BW, Steine EJ, Hanna J, Lodato MA, Frampton GM, Sharp PA, et al. 2010. Histone H3K27ac separates active from poised enhancers and predicts developmental state. Proc Natl Acad Sci USA 107: 21931–21936.

Ding Q, Regan SN, Xia Y, Oostrom LA, Cowan CA, Musunuru K. 2013. Enhanced efficiency of human pluripotent stem cell genome editing through replacing TALENs with CRISPRs. Cell Stem Cell 12: 393–394.

Dobin A, Davis CA, Schlesinger F, Drenkow J, Zaleski C, Jha S, Batut P, Chaisson M, Gingeras TR. 2013. STAR: ultrafast universal RNA-seq aligner. Bioinformatics 29: 15–21.

Domenici G, Aurrekoetxea-Rodríguez I, Simões BM, Rábano M, Lee SY, Millán JS, Comaills V, Oliemuller E, López-Ruiz JA, Zabalza I, et al. 2019. A Sox2–Sox9 signalling axis maintains human breast luminal progenitor and breast cancer stem cells. Oncogene. http://www.nature.com/articles/s41388-018-0656-7 (Accessed January 14, 2019).

Driskell RR, Giangreco A, Jensen KB, Mulder KW, Watt FM. 2009. Sox2-positive dermal papilla cells specify hair follicle type in mammalian epidermis. Development 136: 2815–2823.

Dunn OJ. 1964. Multiple Comparisons Using Rank Sums. Technometrics 6: 241–252.

Dunnett CW. 1955. A Multiple Comparison Procedure for Comparing Several Treatments with a Control. Journal of the American Statistical Association 50: 1096–1121.

ENCODE Project Consortium. 2012. An integrated encyclopedia of DNA elements in the human genome. Nature 489: 57–74.

Fang X, Yoon J-G, Li L, Yu W, Shao J, Hua D, Zheng S, Hood L, Goodlett DR, Foltz G, et al. 2011. The SOX2 response program in glioblastoma multiforme: an integrated ChIP-seq, expression microarray, and microRNA analysis. BMC Genomics 12: 11.

Favaro R, Valotta M, Ferri ALM, Latorre E, Mariani J, Giachino C, Lancini C, Tosetti V, Ottolenghi S, Taylor V, et al. 2009. Hippocampal development and neural stem cell maintenance require Sox2-dependent regulation of Shh. Nature Neuroscience 12: 1248–1256.

Ferone G, Song J-Y, Sutherland KD, Bhaskaran R, Monkhorst K, Lambooij J-P, Proost N, Gargiulo G, Berns A. 2016. SOX2 Is the Determining Oncogenic Switch in Promoting Lung Squamous Cell Carcinoma from Different Cells of Origin. Cancer Cell 30: 519–532.

Ferri ALM, Cavallaro M, Braida D, Di Cristofano A, Canta A, Vezzani A, Ottolenghi S, Pandolfi PP, Sala M, DeBiasi S, et al. 2004. Sox2 deficiency causes neurodegeneration and impaired neurogenesis in the adult mouse brain. Development 131: 3805–3819.

Francis R, Guo H, Streutker C, Ahmed M, Yung T, Dirks PB, He HH, Kim T-H. 2019. Gastrointestinal transcription factors drive lineage-specific developmental programs in organ specification and cancer. Sci Adv 5: eaax8898.

Fu X, Jeselsohn R, Pereira R, Hollingsworth EF, Creighton CJ, Li F, Shea M, Nardone A, Angelis CD, Heiser LM, et al. 2016.FOXA1 overexpression mediates endocrine resistance by altering the ER transcriptome and IL-8 expression in ER-positive breast cancer. PNAS 113: E6600–E6609.

Fu X, Pereira R, De Angelis C, Veeraraghavan J, Nanda S, Qin L, Cataldo ML, Sethunath V, Mehravaran S, Gutierrez C, et al. 2019. FOXA1 upregulation promotes enhancer and transcriptional reprogramming in endocrine-resistant breast cancer. Proc Natl Acad Sci USA 116: 26823–26834.

Fullwood MJ, Liu MH, Pan YF, Liu J, Xu H, Mohamed YB, Orlov YL, Velkov S, Ho A, Mei PH, et al. 2009. An oestrogen-receptor-alpha-bound human chromatin interactome. Nature 462: 58–64.

Gangemi RMR, Griffero F, Marubbi D, Perera M, Capra MC, Malatesta P, Ravetti GL, Zona GL, Daga A, Corte G. 2009. SOX2 silencing in glioblastoma tumor-initiating cells causes stop of proliferation and loss of tumorigenicity. Stem Cells 27: 40–48.

Gertsenstein M, Nutter LMJ. 2021. Production of knockout mouse lines with Cas9. Methods 191: 32–43.

Gontan C, de Munck A, Vermeij M, Grosveld F, Tibboel D, Rottier R. 2008. Sox2 is important for two crucial processes in lung development: Branching morphogenesis and epithelial cell differentiation. Developmental Biology 317: 296–309.

Gopi LK, Kidder BL. 2021. Integrative pan cancer analysis reveals epigenomic variation in cancer type and cell specific chromatin domains. Nat Commun 12: 1419.

Graham V, Khudyakov J, Ellis P, Pevny L. 2003. SOX2 Functions to Maintain Neural Progenitor Identity. Neuron 39: 749–765.

Gründer A, Ebel TT, Mallo M, Schwarzkopf G, Shimizu T, Sippel AE, Schrewe H. 2002. Nuclear factor I-B (Nfib) deficient mice have severe lung hypoplasia. Mech Dev 112: 69–77.

Gupta PB, Fillmore CM, Jiang G, Shapira SD, Tao K, Kuperwasser C, Lander ES. 2011. Stochastic state transitions give rise to phenotypic equilibrium in populations of cancer cells. Cell 146: 633–644.

Hägerstrand D, He X, Bradic Lindh M, Hoefs S, Hesselager G, Ostman A, Nistér M. 2011. Identification of a SOX2-dependent subset of tumor- and sphere-forming glioblastoma cells with a distinct tyrosine kinase inhibitor sensitivity profile. Neuro Oncol 13: 1178–1191.

Harris L, Genovesi LA, Gronostajski RM, Wainwright BJ, Piper M. 2015. Nuclear factor one transcription factors: Divergent functions in developmental versus adult stem cell populations. Dev Dyn 244: 227–238.

Hawkins RD, Hon GC, Lee LK, Ngo Q, Lister R, Pelizzola M, Edsall LE, Kuan S, Luu Y, Klugman S, et al. 2010. Distinct epigenomic landscapes of pluripotent and lineage-committed human cells. Cell Stem Cell 6: 479–491.

Heintzman ND, Hon GC, Hawkins RD, Kheradpour P, Stark A, Harp LF, Ye Z, Lee LK, Stuart RK, Ching CW, et al. 2009. Histone modifications at human enhancers reflect global cell-type-specific gene expression. Nature 459: 108–112.

Heintzman ND, Stuart RK, Hon G, Fu Y, Ching CW, Hawkins RD, Barrera LO, Van Calcar S, Qu C, Ching KA, et al. 2007. Distinct and predictive chromatin signatures of transcriptional promoters and enhancers in the human genome. Nat Genet 39: 311–318.

Heinz S, Benner C, Spann N, Bertolino E, Lin YC, Laslo P, Cheng JX, Murre C, Singh H, Glass CK. 2010. Simple combinations of lineage-determining transcription factors prime cis-regulatory elements required for macrophage and B cell identities. Mol Cell 38: 576–589.

Hnisz D, Abraham BJ, Lee TI, Lau A, Saint-André V, Sigova AA, Hoke HA, Young RA. 2013. Super-Enhancers in the Control of Cell Identity and Disease. Cell 155: 934– 947.

Hoadley KA, Yau C, Wolf DM, Cherniack AD, Tamborero D, Ng S, Leiserson MDM, Niu B, McLellan MD, Uzunangelov V, et al. 2014. Multiplatform analysis of 12 cancer types reveals molecular classification within and across tissues of origin. Cell 158: 929–944.

Holm S. 1979. A Simple Sequentially Rejective Multiple Test Procedure. Scandinavian Journal of Statistics 6: 65–70.

Hsu Y-C, Osinski J, Campbell CE, Litwack ED, Wang D, Liu S, Bachurski CJ, Gronostajski RM. 2011. Mesenchymal nuclear factor I B regulates cell proliferation and epithelial differentiation during lung maturation. Dev Biol 354: 242–252.

Hussenet T, Dali S, Exinger J, Monga B, Jost B, Dembelé D, Martinet N, Thibault C, Huelsken J, Brambilla E, et al. 2010. SOX2 Is an Oncogene Activated by Recurrent 3q26.3 Amplifications in Human Lung Squamous Cell Carcinomas. PLOS ONE 5: e8960.

Iglesias JM, Leis O, Pérez Ruiz E, Gumuzio Barrie J, Garcia-Garcia F, Aduriz A, Beloqui I, Hernandez-Garcia S, Lopez-Mato MP, Dopazo J, et al. 2014. The Activation of the Sox2 RR2 Pluripotency Transcriptional Reporter in Human Breast Cancer Cell Lines is Dynamic and Labels Cells with Higher Tumorigenic Potential. Front Oncol 4: 308.

Jeon H-M, Sohn Y-W, Oh S-Y, Oh S-Y, Kim S-H, Beck S, Kim S, Kim H. 2011. ID4 imparts chemoresistance and cancer stemness to glioma cells by derepressing miR-9*-mediated suppression of SOX2. Cancer Res 71: 3410–3421.

Jeselsohn R, Cornwell M, Pun M, Buchwalter G, Nguyen M, Bango C, Huang Y, Kuang Y, Paweletz C, Fu X, et al. 2017. Embryonic transcription factor SOX9 drives breast cancer endocrine resistance. Proceedings of the National Academy of Sciences 114: E4482–E4491.

Jung K, Gupta N, Wang P, Lewis JT, Gopal K, Wu F, Ye X, Alshareef A, Abdulkarim BS, Douglas DN, et al. 2015. Triple negative breast cancers comprise a highly tumorigenic cell subpopulation detectable by its high responsiveness to a Sox2 regulatory region 2 (SRR2) reporter. Oncotarget 6: 10366–10373.

Kaplan EL, Meier P. 1958. Nonparametric Estimation from Incomplete Observations. Journal of the American Statistical Association 53: 457–481.

Kent WJ, Sugnet CW, Furey TS, Roskin KM, Pringle TH, Zahler AM, Haussler and D. 2002. The Human Genome Browser at UCSC. Genome Res 12: 996–1006.

Kiernan AE, Pelling AL, Leung KKH, Tang ASP, Bell DM, Tease C, Lovell-Badge R, Steel KP, Cheah KSE. 2005. Sox2 is required for sensory organ development in the mammalian inner ear. Nature 434: 1031–1035.

Kılıç Y, Çelebiler AÇ, Sakızlı M. 2014. Selecting housekeeping genes as references for the normalization of quantitative PCR data in breast cancer. Clinical and Translational Oncology 16: 184–190.

Krasnov GS, Kudryavtseva AV, Snezhkina AV, Lakunina VA, Beniaminov AD, Melnikova NV, Dmitriev AA. 2019. Pan-Cancer Analysis of TCGA Data Revealed Promising Reference Genes for qPCR Normalization. Frontiers in Genetics 10. https://www.frontiersin.org/article/10.3389/fgene.2019.00097/full (Accessed March 10, 2020).

Lambert SA, Jolma A, Campitelli LF, Das PK, Yin Y, Albu M, Chen X, Taipale J, Hughes TR, Weirauch MT. 2018. The Human Transcription Factors. Cell 172: 650–665.

Lapidot T, Sirard C, Vormoor J, Murdoch B, Hoang T, Caceres-Cortes J, Minden M, Paterson B, Caligiuri MA, Dick JE. 1994. A cell initiating human acute myeloid leukaemia after transplantation into SCID mice. Nature 367: 645–648.

Leis O, Eguiara A, Lopez-Arribillaga E, Alberdi MJ, Hernandez-Garcia S, Elorriaga K, Pandiella A, Rezola R, Martin AG. 2012. Sox2 expression in breast tumours and activation in breast cancer stem cells. Oncogene 31: 1354–1365.

Li L, Wu D, Yu Q, Li L, Wu P. 2017. Prognostic value of FOXM1 in solid tumors: a systematic review and meta-analysis. Oncotarget 8: 32298–32308.

Li Y, Rivera CM, Ishii H, Jin F, Selvaraj S, Lee AY, Dixon JR, Ren B. 2014. CRISPR Reveals a Distal Super-Enhancer Required for Sox2 Expression in Mouse Embryonic Stem Cells ed. Q. Wu. PLoS ONE 9: e114485.

Liang S, Furuhashi M, Nakane R, Nakazawa S, Goudarzi H, Hamada J, Iizasa H. 2013. Isolation and characterization of human breast cancer cells with SOX2 promoter activity. Biochemical and Biophysical Research Communications 437: 205–211.

Liao Y, Smyth GK, Shi W. 2014. featureCounts: an efficient general purpose program for assigning sequence reads to genomic features. Bioinformatics 30: 923–930.

Ling G-Q, Chen, Dong-bo, Wang, Bao-Qing, Zhang, Lan-Sheng. 2012. Expression of the pluripotency markers Oct3/4, Nanog and Sox2 in human breast cancer cell lines. Oncology Letters. http://www.spandidos-publications.com/10.3892/ol.2012.916 (Accessed November 9, 2016).

Liu C, Wang M, Wei X, Wu L, Xu J, Dai X, Xia J, Cheng M, Yuan Y, Zhang P, et al. 2019. An ATAC-seq atlas of chromatin accessibility in mouse tissues. Sci Data 6: 65.

Liu J, Lichtenberg T, Hoadley KA, Poisson LM, Lazar AJ, Cherniack AD, Kovatich AJ, Benz CC, Levine DA, Lee AV, et al. 2018a. An Integrated TCGA Pan-Cancer Clinical Data Resource to Drive High-Quality Survival Outcome Analytics. Cell 173: 400–416.e11.

Liu K-C, Lin B-S, Zhao M, Yang X, Chen M, Gao A, Que J, Lan X-P. 2013. The multiple roles for Sox2 in stem cell maintenance and tumorigenesis. Cell Signal 25. http://www.ncbi.nlm.nih.gov/pmc/articles/PMC3871517/ (Accessed April 5, 2017).

Liu P, Tang H, Song C, Wang J, Chen B, Huang X, Pei X, Liu L. 2018b. SOX2 Promotes Cell Proliferation and Metastasis in Triple Negative Breast Cancer. Front Pharmacol 9. https://www.ncbi.nlm.nih.gov/pmc/articles/PMC6110877/ (Accessed March 21, 2019).

Liu Y, Wu Z, Zhou J, Ramadurai DKA, Mortenson KL, Aguilera-Jimenez E, Yan Y, Yang X, Taylor AM, Varley KE, et al. 2021. A predominant enhancer co-amplified with the SOX2 oncogene is necessary and sufficient for its expression in squamous cancer. Nat Commun 12: 7139.

Liu Y, Zhao Y, Skerry B, Wang X, Colin-Cassin C, Radisky DC, Kaestner KH, Li Z. 2016. Foxa1 is essential for mammary duct formation. Genesis 54: 277–285.

Love MI, Huber W, Anders S. 2014. Moderated estimation of fold change and dispersion for RNA-seq data with DESeq2. Genome Biol 15. https://www.ncbi.nlm.nih.gov/pmc/articles/PMC4302049/ (Accessed November 18, 2017).

Lovén J, Hoke HA, Lin CY, Lau A, Orlando DA, Vakoc CR, Bradner JE, Lee TI, Young RA. 2013. Selective Inhibition of Tumor Oncogenes by Disruption of Super-Enhancers. Cell 153: 320–334.

Lupien M, Eeckhoute J, Meyer CA, Wang Q, Zhang Y, Li W, Carroll JS, Liu XS, Brown M. 2008. FoxA1 Translates Epigenetic Signatures into Enhancer-Driven Lineage-Specific Transcription. Cell 132: 958–970.

Lyng MB, Lænkholm A-V, Pallisgaard N, Ditzel HJ. 2008. Identification of genes for normalization of real-time RT-PCR data in breast carcinomas. BMC Cancer 8. http://bmccancer.biomedcentral.com/articles/10.1186/1471-2407-8-20 (Accessed June 17, 2020).

Maier S, Wilbertz T, Braun M, Scheble V, Reischl M, Mikut R, Menon R, Nikolov P, Petersen K, Beschorner C, et al. 2011. SOX2 amplification is a common event in squamous cell carcinomas of different organ sites. Human Pathology 42: 1078– 1088.

Mali P, Yang L, Esvelt KM, Aach J, Guell M, DiCarlo JE, Norville JE, Church GM. 2013. RNA-Guided Human Genome Engineering via Cas9. Science 339: 823–826.

Meng Y, Xu Q, Chen L, Wang L, Hu X. 2020. The function of SOX2 in breast cancer and relevant signaling pathway. Pathology - Research and Practice 216: 153023.

Meuleman W, Muratov A, Rynes E, Halow J, Lee K, Bates D, Diegel M, Dunn D, Neri F, Teodosiadis A, et al. 2020. Index and biological spectrum of human DNase I hypersensitive sites. Nature 584: 244–251.

Minoo P, Su G, Drum H, Bringas P, Kimura S. 1999. Defects in tracheoesophageal and lung morphogenesis in Nkx2.1(-/-) mouse embryos. Dev Biol 209: 60–71.

Mitchell JA, Clay I, Umlauf D, Chen C-Y, Moir CA, Eskiw CH, Schoenfelder S, Chakalova L, Nagano T, Fraser P. 2012. Nuclear RNA sequencing of the mouse erythroid cell transcriptome. PLoS ONE 7: e49274.

Miyagi S, Nishimoto M, Saito T, Ninomiya M, Sawamoto K, Okano H, Muramatsu M, Oguro H, Iwama A, Okuda A. 2006. The Sox2 Regulatory Region 2 Functions as a Neural Stem Cell-specific Enhancer in the Telencephalon. Journal of Biological Chemistry 281: 13374–13381.

Miyagi S, Saito T, Mizutani K, Masuyama N, Gotoh Y, Iwama A, Nakauchi H, Masui S, Niwa H, Nishimoto M, et al. 2004. The Sox-2 regulatory regions display their activities in two distinct types of multipotent stem cells. Mol Cell Biol 24: 4207– 4220.

Moorthy SD, Davidson S, Shchuka VM, Singh G, Malek-Gilani N, Langroudi L, Martchenko A, So V, Macpherson NN, Mitchell JA. 2017. Enhancers and super-enhancers have an equivalent regulatory role in embryonic stem cells through regulation of single or multiple genes. Genome Res 27: 246–258.

Nakatsugawa M, Takahashi A, Hirohashi Y, Torigoe T, Inoda S, Murase M, Asanuma H, Tamura Y, Morita R, Michifuri Y, et al. 2011. SOX2 is overexpressed in stem-like cells of human lung adenocarcinoma and augments the tumorigenicity. Lab Invest 91: 1796–1804.

Novak D, Hüser L, Elton JJ, Umansky V, Altevogt P, Utikal J. 2019. SOX2 in development and cancer biology. Seminars in Cancer Biology. http://www.sciencedirect.com/science/article/pii/S1044579X18301858 (Accessed August 19, 2019).

Okabe A, Kaneda A. 2021. Transcriptional dysregulation by aberrant enhancer activation and rewiring in cancer. Cancer Sci 112: 2081–2088.

Okubo T, Pevny LH, Hogan BLM. 2006. Sox2 is required for development of taste bud sensory cells. Genes & Development 20: 2654–2659.

Orstad G, Fort G, Parnell TJ, Jones A, Stubben C, Lohman B, Gillis KL, Orellana W, Tariq R, Klingbeil O, et al. 2022. FoxA1 and FoxA2 control growth and cellular identity in NKX2-1-positive lung adenocarcinoma. Dev Cell 57: 1866–1882.e10.

Otsubo T, Akiyama Y, Yanagihara K, Yuasa Y. 2008. SOX2 is frequently downregulated in gastric cancers and inhibits cell growth through cell-cycle arrest and apoptosis. Br J Cancer 98: 824–831.

Paranjapye A, Mutolo MJ, Ebron JS, Leir S-H, Harris A. 2020. The FOXA1 transcriptional network coordinates key functions of primary human airway epithelial cells. Am J Physiol Lung Cell Mol Physiol 319: L126–L136.

Pennacchio LA, Ahituv N, Moses AM, Prabhakar S, Nobrega MA, Shoukry M, Minovitsky S, Dubchak I, Holt A, Lewis KD, et al. 2006. In vivo enhancer analysis of human conserved non-coding sequences. Nature 444: 499–502.

Pierrou S, Hellqvist M, Samuelsson L, Enerbäck S, Carlsson P. 1994. Cloning and characterization of seven human forkhead proteins: binding site specificity and DNA bending. The EMBO Journal 13: 5002–5012.

Piva M, Domenici G, Iriondo O, Rábano M, Simões BM, Comaills V, Barredo I, López-Ruiz JA, Zabalza I, Kypta R, et al. 2014. Sox2 promotes tamoxifen resistance in breast cancer cells. EMBO Molecular Medicine 6: 66–79.

Pollard KS, Hubisz MJ, Rosenbloom KR, Siepel A. 2010. Detection of nonneutral substitution rates on mammalian phylogenies. Genome Res 20: 110–121.

Pomerantz MM, Li F, Takeda DY, Lenci R, Chonkar A, Chabot M, Cejas P, Vazquez F, Cook J, Shivdasani RA, et al. 2015. The androgen receptor cistrome is extensively reprogrammed in human prostate tumorigenesis. Nat Genet 47: 1346–1351.

Que J, Choi M, Ziel JW, Klingensmith J, Hogan BLM. 2006. Morphogenesis of the trachea and esophagus: current players and new roles for noggin and Bmps. Differentiation 74: 422–437.

Que J, Luo X, Schwartz RJ, Hogan BLM. 2009. Multiple roles for Sox2 in the developing and adult mouse trachea. Development 136: 1899–1907.

Que J, Okubo T, Goldenring JR, Nam K-T, Kurotani R, Morrisey EE, Taranova O, Pevny LH, Hogan BLM. 2007. Multiple dose-dependent roles for Sox2 in the patterning and differentiation of anterior foregut endoderm. Development 134: 2521–2531.

Rada-Iglesias A, Bajpai R, Swigut T, Brugmann SA, Flynn RA, Wysocka J. 2011. A unique chromatin signature uncovers early developmental enhancers in humans. Nature 470: 279–283.

Ran FA, Hsu PD, Wright J, Agarwala V, Scott DA, Zhang F. 2013. Genome engineering using the CRISPR-Cas9 system. Nat Protoc 8: 2281–2308.

Richart L, Bidard F-C, Margueron R. 2021. Enhancer rewiring in tumors: an opportunity for therapeutic intervention. Oncogene 40: 3475–3491.

Roadmap Epigenomics Consortium, Kundaje A, Meuleman W, Ernst J, Bilenky M, Yen A, Heravi-Moussavi A, Kheradpour P, Zhang Z, Wang J, et al. 2015. Integrative analysis of 111 reference human epigenomes. Nature 518: 317–330.

Roe J-S, Hwang C-I, Somerville TDD, Milazzo JP, Lee EJ, Da Silva B, Maiorino L, Tiriac H, Young CM, Miyabayashi K, et al. 2017. Enhancer Reprogramming Promotes Pancreatic Cancer Metastasis. Cell 170: 875–888.e20.

Rubin AJ, Barajas BC, Furlan-Magaril M, Lopez-Pajares V, Mumbach MR, Howard I, Kim DS, Boxer LD, Cairns J, Spivakov M, et al. 2017. Lineage-specific dynamic and pre-established enhancer-promoter contacts cooperate in terminal differentiation. Nat Genet 49: 1522–1528.

Saenz-Antoñanzas A, Moncho-Amor V, Auzmendi-Iriarte J, Elua-Pinin A, Rizzoti K, Lovell-Badge R, Matheu A. 2021. CRISPR/Cas9 Deletion of SOX2 Regulatory Region 2 (SRR2) Decreases SOX2 Malignant Activity in Glioblastoma. Cancers (Basel) 13: 1574.

Sato T, Yoo S, Kong R, Sinha A, Chandramani-Shivalingappa P, Patel A, Fridrikh M, Nagano O, Masuko T, Beasley MB, et al. 2019. Epigenomic Profiling Discovers Trans-lineage SOX2 Partnerships Driving Tumor Heterogeneity in Lung Squamous Cell Carcinoma. Cancer Research 79: 6084–6100.

Seibt KM, Schmidt T, Heitkam T. 2018. FlexiDot: highly customizable, ambiguity-aware dotplots for visual sequence analyses ed. J. Hancock. Bioinformatics 34: 3575–3577.

Shen L, Shao N, Liu X, Nestler E. 2014. ngs.plot: Quick mining and visualization of next-generation sequencing data by integrating genomic databases. BMC Genomics 15: 284.

Sholl LM, Barletta JA, Yeap BY, Chirieac LR, Hornick JL. 2010. Sox2 protein expression is an independent poor prognostic indicator in stage I lung adenocarcinoma. Am J Surg Pathol 34: 1193–1198.

Sievers F, Wilm A, Dineen D, Gibson TJ, Karplus K, Li W, Lopez R, McWilliam H, Remmert M, Söding J, et al. 2011. Fast, scalable generation of high-quality protein multiple sequence alignments using Clustal Omega. Mol Syst Biol 7: 539.

Simões BM, Piva M, Iriondo O, Comaills V, López-Ruiz JA, Zabalza I, Mieza JA, Acinas O, Vivanco M d.M. 2011. Effects of estrogen on the proportion of stem cells in the breast. Breast Cancer Research and Treatment 129: 23–35.

Singh G, Mullany S, Moorthy SD, Zhang R, Mehdi T, Tian R, Duncan AG, Moses AM, Mitchell JA. 2021. A flexible repertoire of transcription factor binding sites and a diversity threshold determines enhancer activity in embryonic stem cells. Genome Res 31: 564–575.

Singh S, Trevino J, Bora-Singhal N, Coppola D, Haura E, Altiok S, Chellappan SP. 2012. EGFR/Src/Akt signaling modulates Sox2 expression and self-renewal of stem-like side-population cells in non-small cell lung cancer. Molecular Cancer 11: 73.

Soufi A, Donahue G, Zaret KS. 2012. Facilitators and Impediments of the Pluripotency Reprogramming Factors’ Initial Engagement with the Genome. Cell 151: 994– 1004.

Soufi A, Garcia MF, Jaroszewicz A, Osman N, Pellegrini M, Zaret KS. 2015. Pioneer transcription factors target partial DNA motifs on nucleosomes to initiate reprogramming. Cell 161: 555–568.

Soule HD, Vazquez J, Long A, Albert S, Brennan M. 1973. A human cell line from a pleural effusion derived from a breast carcinoma. Journal of the National Cancer Institute 51: 1409–1416.

Stark R, Brown G. 2011. DiffBind: differential binding analysis of ChIP-Seq peak data. http://bioconductor.org/packages/release/bioc/vignettes/DiffBind/inst/doc/DiffBind.pdf.

Steele-Perkins G, Plachez C, Butz KG, Yang G, Bachurski CJ, Kinsman SL, Litwack ED, Richards LJ, Gronostajski RM. 2005. The transcription factor gene Nfib is essential for both lung maturation and brain development. Mol Cell Biol 25: 685–698.

Stergachis AB, Neph S, Reynolds A, Humbert R, Miller B, Paige SL, Vernot B, Cheng JB, Thurman RE, Sandstrom R, et al. 2013. Developmental Fate and Cellular Maturity Encoded in Human Regulatory DNA Landscapes. Cell 154: 888–903.

Stolzenburg S, Rots MG, Beltran AS, Rivenbark AG, Yuan X, Qian H, Strahl BD, Blancafort P. 2012. Targeted silencing of the oncogenic transcription factor SOX2 in breast cancer. Nucleic Acids Research 40: 6725–6740.

Sun C, Sun L, Li Y, Kang X, Zhang S, Liu Y. 2013. Sox2 expression predicts poor survival of hepatocellular carcinoma patients and it promotes liver cancer cell invasion by activating Slug. Med Oncol 30: 503.

Takahashi K, Tanabe K, Ohnuki M, Narita M, Ichisaka T, Tomoda K, Yamanaka S. 2007. Induction of Pluripotent Stem Cells from Adult Human Fibroblasts by Defined Factors. Cell 131: 861–872.

Takahashi K, Yamanaka S. 2006. Induction of Pluripotent Stem Cells from Mouse Embryonic and Adult Fibroblast Cultures by Defined Factors. Cell 126: 663–676.

Takeda K, Mizushima T, Yokoyama Y, Hirose H, Wu X, Qian Y, Ikehata K, Miyoshi N, Takahashi H, Haraguchi N, et al. 2018. Sox2 is associated with cancer stem-like properties in colorectal cancer. Scientific Reports 8: 17639.

Talebi A, Kianersi K, Beiraghdar M. 2015. Comparison of gene expression of SOX2 and OCT4 in normal tissue, polyps, and colon adenocarcinoma using immunohistochemical staining. Adv Biomed Res 4: 234.

Taranova OV, Magness ST, Fagan BM, Wu Y, Surzenko N, Hutton SR, Pevny LH. 2006. SOX2 is a dose-dependent regulator of retinal neural progenitor competence. Genes Dev 20: 1187–1202.

Teramoto M, Sugawara R, Minegishi K, Uchikawa M, Takemoto T, Kuroiwa A, Ishii Y, Kondoh H. 2020. The absence of SOX2 in the anterior foregut alters the esophagus into trachea and bronchi in both epithelial and mesenchymal components. Biology Open 9: bio048728.

Thurman RE, Rynes E, Humbert R, Vierstra J, Maurano MT, Haugen E, Sheffield NC, Stergachis AB, Wang H, Vernot B, et al. 2012. The accessible chromatin landscape of the human genome. Nature 489: 75–82.

Tobias IC, Abatti LE, Moorthy SD, Mullany S, Taylor T, Khader N, Filice MA, Mitchell JA. 2021. Transcriptional enhancers: from prediction to functional assessment on a genome-wide scale. Genome 64: 426–448.

Tolhuis B, Palstra R-J, Splinter E, Grosveld F, de Laat W. 2002. Looping and Interaction between Hypersensitive Sites in the Active β-globin Locus. Molecular Cell 10: 1453–1465.

Tomioka M, Nishimoto M, Miyagi S, Katayanagi T, Fukui N, Niwa H, Muramatsu M, Okuda A. 2002. Identification of Sox-2 regulatory region which is under the control of Oct-3/4–Sox-2 complex. Nucleic Acids Res 30: 3202–3213.

Tompkins DH, Besnard V, Lange AW, Keiser AR, Wert SE, Bruno MD, Whitsett JA. 2011. Sox2 Activates Cell Proliferation and Differentiation in the Respiratory Epithelium. Am J Respir Cell Mol Biol 45: 101–110.

Tompkins DH, Besnard V, Lange AW, Wert SE, Keiser AR, Smith AN, Lang R, Whitsett JA. 2009. Sox2 Is Required for Maintenance and Differentiation of Bronchiolar Clara, Ciliated, and Goblet Cells ed. C.M. Verfaillie. PLoS ONE 4: e8248.

Tukey JW. 1949. Comparing Individual Means in the Analysis of Variance. Biometrics 5: 99.

Wang S, Tie J, Wang R, Hu F, Gao L, Wang W, Wang L, Li Z, Hu S, Tang S, et al. 2015. SOX2, a predictor of survival in gastric cancer, inhibits cell proliferation and metastasis by regulating PTEN. Cancer Lett 358: 210–219.

Williamson KA, Hever AM, Rainger J, Rogers RC, Magee A, Fiedler Z, Keng WT, Sharkey FH, McGill N, Hill CJ, et al. 2006. Mutations in SOX2 cause anophthalmia-esophageal-genital (AEG) syndrome. Hum Mol Genet 15: 1413–1422.

Woolfe A, Goodson M, Goode DK, Snell P, McEwen GK, Vavouri T, Smith SF, North P, Callaway H, Kelly K, et al. 2005. Highly conserved non-coding sequences are associated with vertebrate development. PLoS Biol 3: e7.

Wu C, Zhu X, Liu W, Ruan T, Wan W, Tao K. 2018. NFIB promotes cell growth, aggressiveness, metastasis and EMT of gastric cancer through the Akt/Stat3 signaling pathway. Oncol Rep 40: 1565–1573.

Wuebben EL, Rizzino A. 2017. The dark side of SOX2: cancer-a comprehensive overview. Oncotarget 8: 44917–44943.

Yu G, Wang L-G, Han Y, He Q-Y. 2012. clusterProfiler: an R Package for Comparing Biological Themes Among Gene Clusters. OMICS: A Journal of Integrative Biology 16: 284–287.

Yu J, Vodyanik MA, Smuga-Otto K, Antosiewicz-Bourget J, Frane JL, Tian S, Nie J, Jonsdottir GA, Ruotti V, Stewart R, et al. 2007. Induced Pluripotent Stem Cell Lines Derived from Human Somatic Cells. Science 318: 1917–1920.

Zappone MV, Galli R, Catena R, Meani N, De Biasi S, Mattei E, Tiveron C, Vescovi AL, Lovell-Badge R, Ottolenghi S, et al. 2000. Sox2 regulatory sequences direct expression of a (beta)-geo transgene to telencephalic neural stem cells and precursors of the mouse embryo, revealing regionalization of gene expression in CNS stem cells. Development 127: 2367–2382.

Zenteno JC, Perez-Cano HJ, Aguinaga M. 2006. Anophthalmia-esophageal atresia syndrome caused by an SOX2 gene deletion in monozygotic twin brothers with markedly discordant phenotypes. Am J Med Genet A 140: 1899–1903.

Zhang J-P, Li X-L, Li G-H, Chen W, Arakaki C, Botimer GD, Baylink D, Zhang L, Wen W, Fu Y-W, et al. 2017. Efficient precise knockin with a double cut HDR donor after CRISPR/Cas9-mediated double-stranded DNA cleavage. Genome Biology 18: 35.

Zhang L-H, Yin Y-H, Chen H-Z, Feng S-Y, Liu J-L, Chen L, Fu W-L, Sun G-C, Yu X-G, Xu D-G. 2020a. TRIM24 promotes stemness and invasiveness of glioblastoma cells via activating Sox2 expression. Neuro Oncol 22.

Zhang X, Yu H, Yang Y, Zhu R, Bai J, Peng Z, He Y, Chen L, Chen W, Fang D, et al. 2010. SOX2 in gastric carcinoma, but not Hath1, is related to patients’ clinicopathological features and prognosis. J Gastrointest Surg 14: 1220–1226.

Zhang X-H, Wang W, Wang Y-Q, Zhu L, Ma L. 2020b. The association of SOX2 with clinical features and prognosis in colorectal cancer: A meta-analysis. Pathol Res Pract 216.

Zhou HY, Katsman Y, Dhaliwal NK, Davidson S, Macpherson NN, Sakthidevi M, Collura F, Mitchell JA. 2014a. A Sox2 distal enhancer cluster regulates embryonic stem cell differentiation potential. Genes & Development 28: 2699–2711.

Zhou M, Zhou L, Zheng L, Guo L, Wang Y, Liu H, Ou C, Ding Z. 2014b. miR-365 promotes cutaneous squamous cell carcinoma (CSCC) through targeting nuclear factor I/B (NFIB). PLoS One 9: e100620.

Zhu J, Adli M, Zou JY, Verstappen G, Coyne M, Zhang X, Durham T, Miri M, Deshpande V, De Jager PL, et al. 2013. Genome-wide chromatin state transitions associated with developmental and environmental cues. Cell 152: 642–654.

Zhu LJ, Gazin C, Lawson ND, Pagès H, Lin SM, Lapointe DS, Green MR. 2010. ChIPpeakAnno: a Bioconductor package to annotate ChIP-seq and ChIP-chip data. BMC Bioinformatics 11. https://bmcbioinformatics.biomedcentral.com/articles/10.1186/1471-2105-11-237 (Accessed June 17, 2020).

Zhu Y, Huang S, Chen S, Chen J, Wang Z, Wang Y, Zheng H. 2021. SOX2 promotes chemoresistance, cancer stem cells properties, and epithelial-mesenchymal transition by β-catenin and Beclin1/autophagy signaling in colorectal cancer. Cell Death Dis 12: 449.

