## Supplementary figures and images for "Epigenetic reprogramming of a distal developmental enhancer cluster drives *SOX2* overexpression in breast and lung cancer"

### Supplemental Figure S1

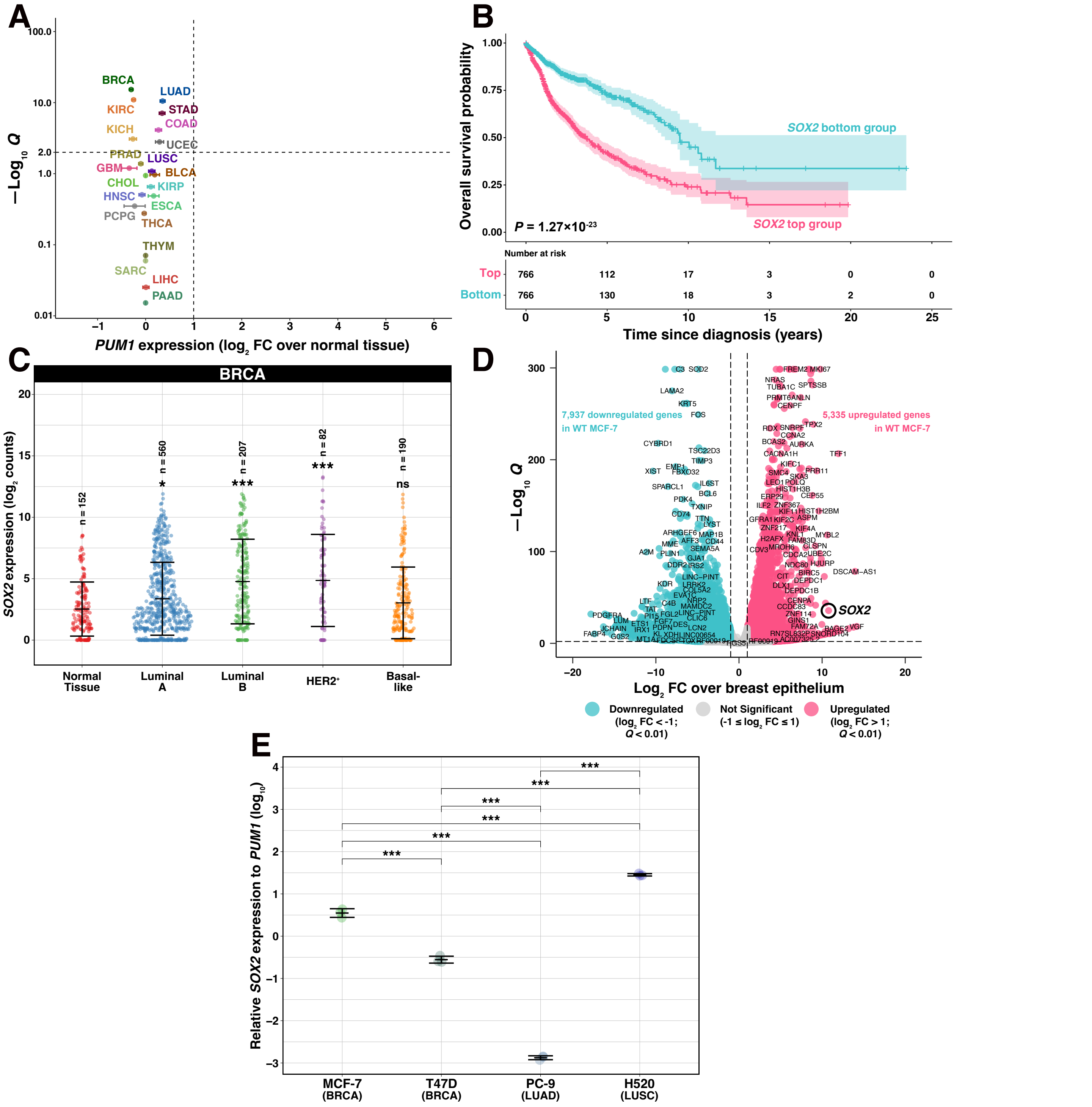

### Supplemental Figure S2

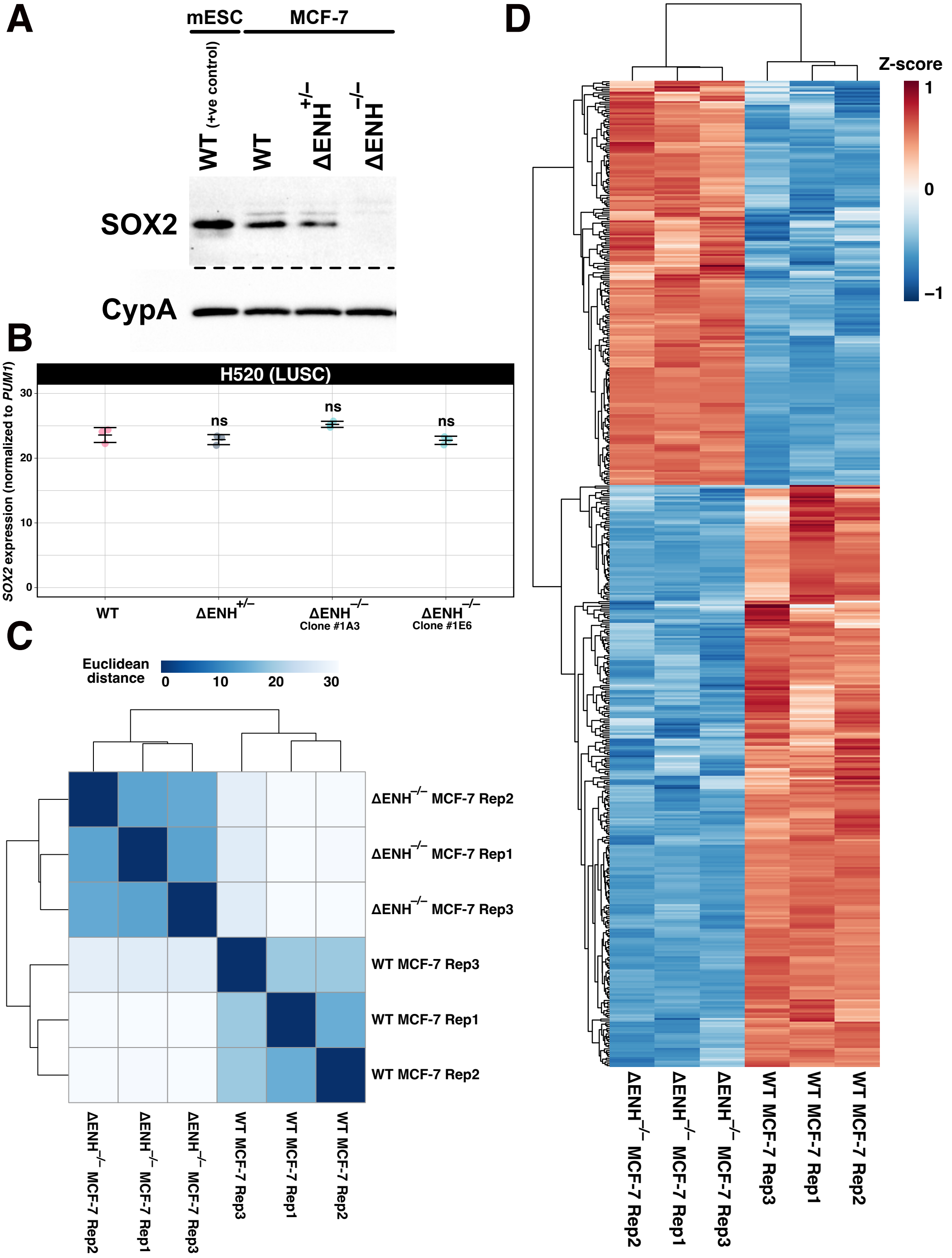

### Supplemental Figure S3

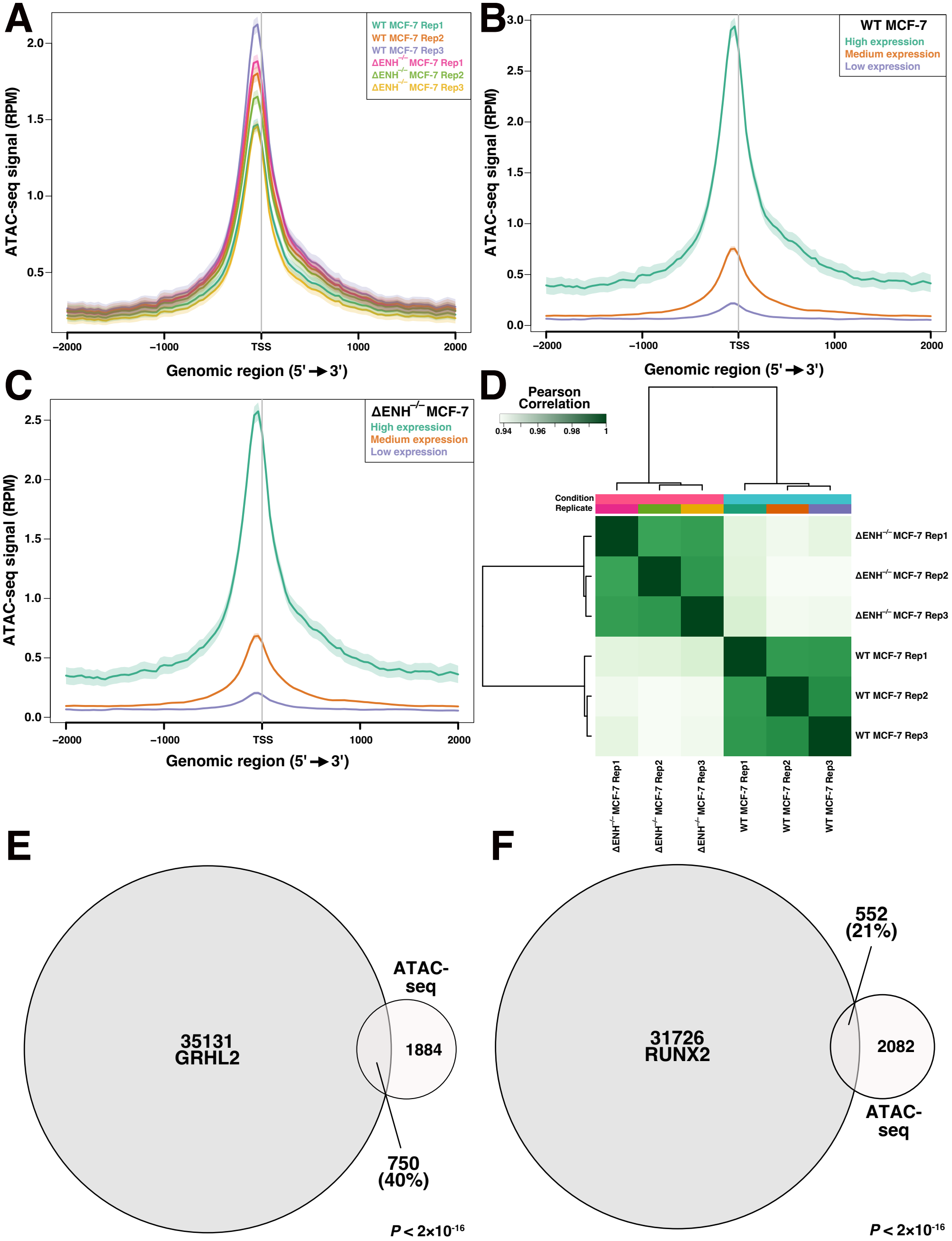

### Supplemental Figure S4

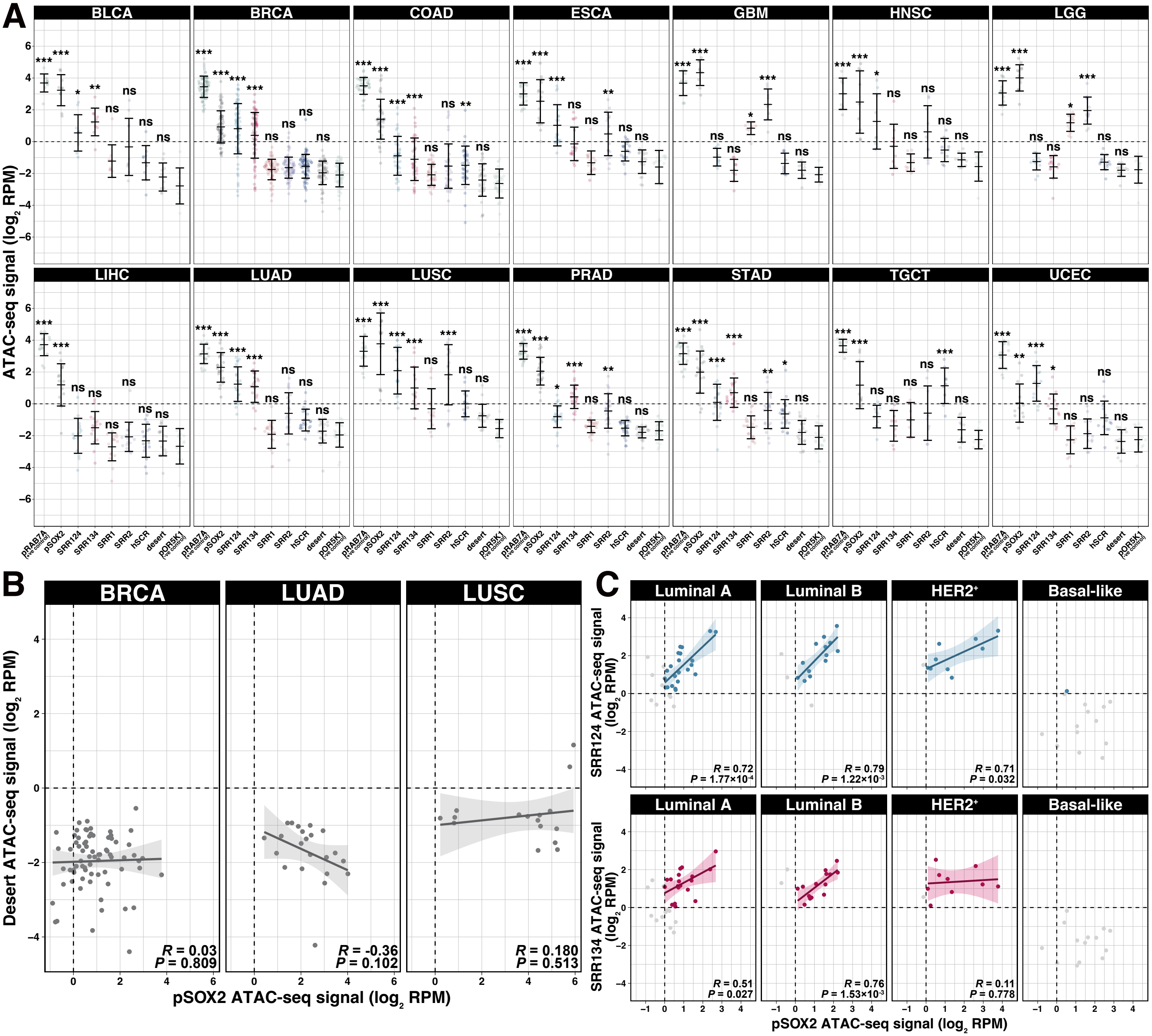

### Supplemental Figure S6

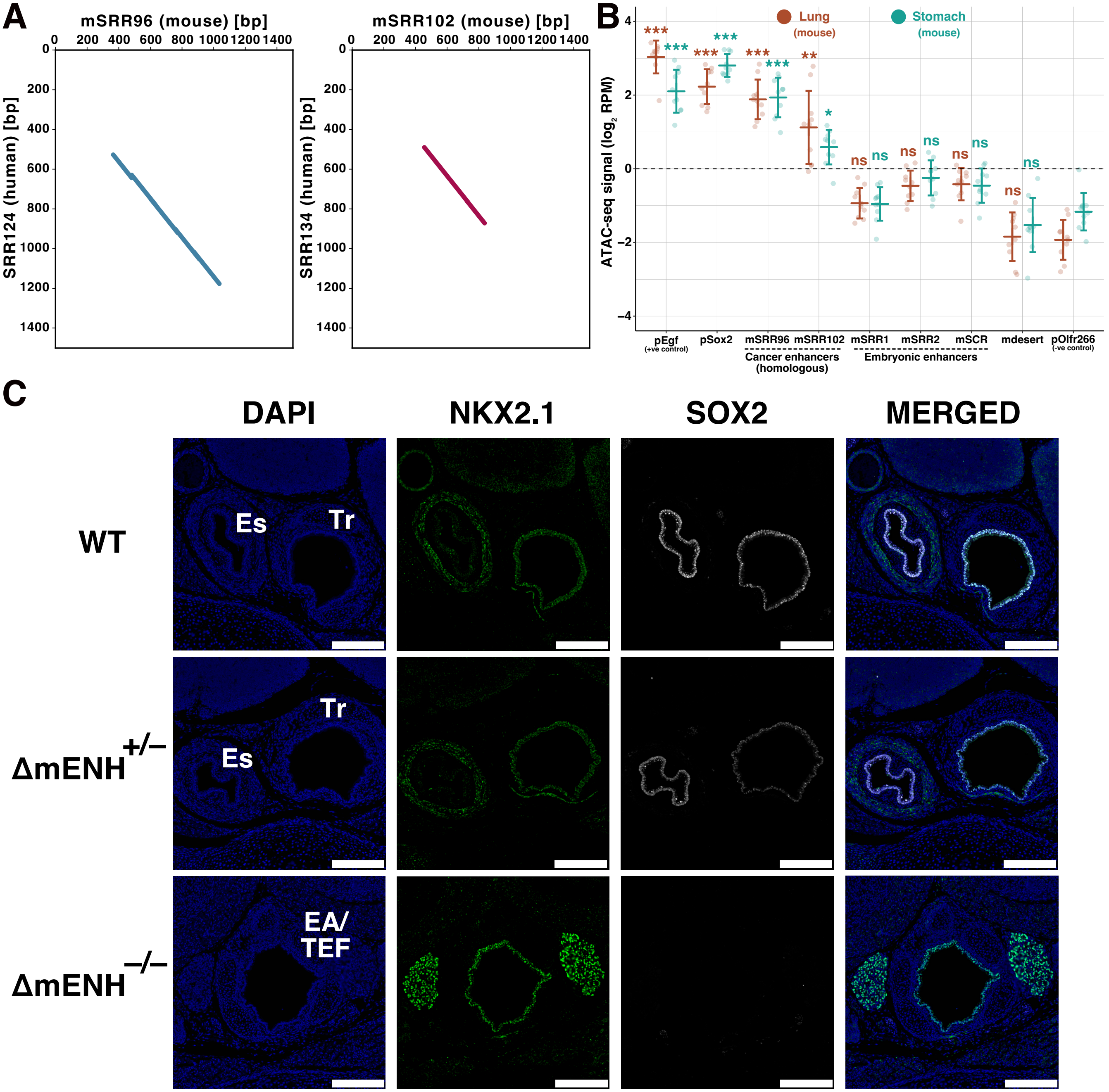
