## Supplemental Figure S5 for "Epigenetic reprogramming of a distal developmental enhancer cluster drives *SOX2* overexpression in breast and lung cancer"

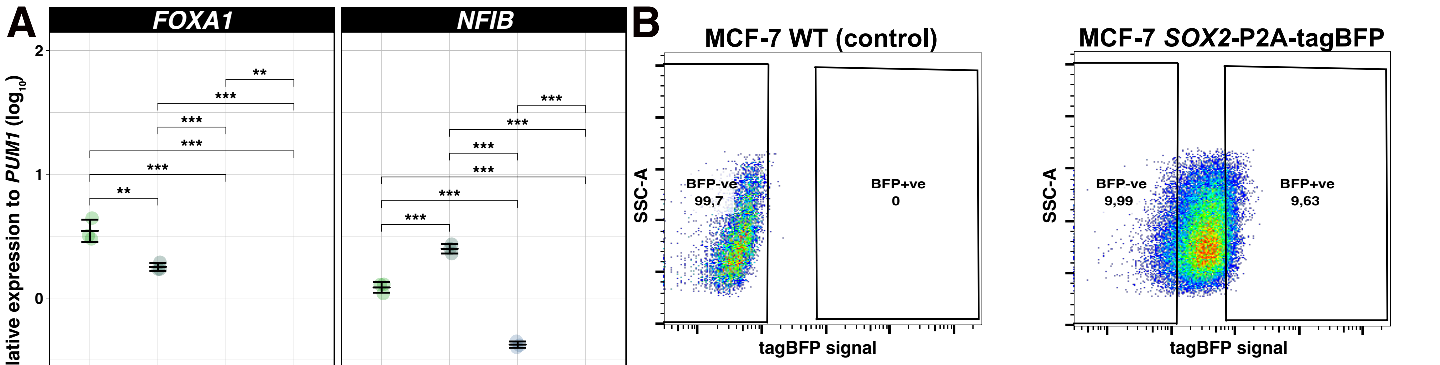


**MCF-7 WT (control)**

| | Genotype | % of population |
|---|---|---|
| ■ | tagBFP negative | 99,7 |
| ■ | tagBFP positive | 0 |

**MCF-7 SOX2-P2A-tagBFP**

| | Genotype | % of population |
|---|---|---|
| ■ | tagBFP negative | 9,99 |
| ■ | tagBFP positive | 9,63 |
